# Differential gene expression drives cell-cycle-dependent transition from monopolar to bipolar growth in fission yeast

**DOI:** 10.1101/2025.08.13.670185

**Authors:** Samridhi Pathak, Maitreyi Das

## Abstract

Cell polarity is important for maintaining cell structure and function. In *S. pomb*e, after division, cells are monopolar and grow from the old end that existed from the previous generation. Cells transition to bipolar growth once the cell reaches a certain size, when the new end generated after division initiates growth. Bipolarity occurs as a consequence of cell growth, where the two cell ends have sufficient growth-promoting factors. However, the transition to bipolarity does not occur in G1-arrested cells even with enhanced cell growth, suggesting a role for the cell-cycle phase in this process. To identify how the cell cycle impacts monopolar to bipolar transition, we performed high-throughput mRNA sequencing to detect differentially expressed genes in G1/S-arrested and G2-phase cells of the cell cycle mutant *cdc10-129*. Gene expression analysis using the DESeq2 R package identified 65 unique genes upregulated in G1/S phase and 35 in G2 phase cells. Enrichment analysis shows that G1/S phase cells upregulated the MAPK pheromone-response pathway, protein folding, rRNA processing, and heat-shock protein binding. G2 phase cells showed upregulation of plasma membrane maintenance and cell wall organization. Protein-protein interaction networks identified the *cdc15*-*hob3*-*rho1*-*bgs1* hub in G2 phase cells. These genes are known to promote bipolar growth and regulate cell wall biogenesis. In G1/S phase cells *spk1-byr2-ste11* hub was identified, which is required for pheromone-response and nutritional stress-dependent G1-arrest. We find that Spk1 also prevents precocious bipolar growth. Thus, we hypothesize that the pheromone response pathway, in addition to promoting G1-G0 transition, also prevents bipolar growth. In nutrient-rich conditions, this pathway is downregulated, G1-arrest is alleviated, the cell cycle progresses, and bipolarity occurs. Our findings suggest that in nutrition-rich conditions, stress response pathways are downregulated enabling transition from monopolar to bipolar growth, along with cell cycle progression.

## Introduction

Cells undergo polarization characterized by the asymmetric delivery and confinement of molecules and biological processes to establish and maintain cell shape [1–3]. Maintenance of cell shape is essential for tissue organization and proper function of the organism [3]. Cell polarity is a result of dynamic interactions of the plasma membrane with cytoskeletal proteins and the subsequent spatiotemporal organization of polarity complexes [2, 4–6]. The molecular nature of the intrinsic and extrinsic spatial cues that establish structural and molecular asymmetry at the cell surface, although highly conserved, is poorly understood [6–8].

From yeast to humans, the key components that establish polarization, such as the Rho GTPases, are conserved, indicating that the underlying mechanisms have not significantly changed throughout evolution [5, 9–13]. The fission yeast, *S. pombe* cells offer a simpler bipolar model of cell polarization regulated via the small GTPases Cdc42, a major regulator of cell polarity [10, 14–17]. Cdc42 is activated by GEFs (guanidine-nucleotide-exchange factors) and inactivated by GAPs (GTPase-activating proteins) [5, 6, 10, 17–19].

In fission yeast, the cell polarity pattern is cell-cycle-dependent. Post cell division, during the early G2 phase, cells are monopolar and initiate growth from the old end that existed in the previous generation. The new end, which is generated during cell division, initiates growth, resulting in bipolar cells after a process called new end take off (NETO). However, what drives NETO during bipolar growth is poorly understood [5, 18]. Previous models suggest that bipolarity occurs once the cell achieves a certain cell size during growth in the G2 phase [5, 18].

In the G2 Phase, oscillations of GTP-bound Cdc42 between the two ends, in an anti-correlated manner, promote bipolar growth [5, 18, 20–22]. The anti-correlated oscillations are due to the competition between the two growing ends for Cdc42 activation [5, 18, 21]. As the cells increase in size and synthesize sufficient proteins, the competition between the ends is reduced [8, 18, 23, 24]. This suggested that as the cell size increased, bipolar Cdc42 activation at the cell ends is efficiently established, allowing simultaneous growth at these sites. A second model defines that Gef1, a secondary activator of Cdc42, acts as a spark-plug to initiate the Cdc42 positive feedback at the new end to allow bipolar growth [20, 22].

It has also been shown that the transition to bipolar growth is cell-cycle-dependent. Cell-cycle mutants arrested in the G1/S phase show increased cell size but remain monopolar [25]. This indicates that bipolarity occurs in the G2 phase, and that cell size increase alone does not explain the transition from monopolar to bipolar growth. In *cdc10-129* mutants under restrictive conditions, the cells arrest in G1/S and remain monopolar [25–27]. Cdc10 protein is a DNA-binding transcription factor responsible for transcription of genes such as *cdt2*, *cig2*, cdc18, *mik1*, *nrm1*, *rec11*, *rec8*, and *yox1* [27, 28]. In the *cdc10-129* temperature-sensitive mutants, the cells grow in length but remain monopolar under restrictive conditions [26–28].

Research in cell polarization mainly examines the Rho GTPase signaling pathways and their downstream effects at the polarized sites. Changes in the regulation of the signaling pathways result in phenotypes such as loss of polarity or monopolarity with corresponding changes in cell dimensions [10, 18, 21, 22, 29–32]. However, the growing ends of the monopolar *cdc10-129* mutant do not show any major difference in the regulation of Cdc42 or cellular dimensions. This suggests that the monopolarity defect observed in these cells could be due to a defect in the global regulation of polarity. We asked if the transition from monopolar to bipolar growth is impacted by the differences in gene expression in the different cell-cycle stages. Indeed, gene expression data from different organisms suggest that cell polarity is also regulated at the systems level. Polarity genes show divergent expression profiles in distinct human tissues and cancer cells [33]. *Drosophila* embryos exhibit cell-cycle-dependent changes in polarity gene expression [34]. Nevertheless, it remains unclear how changes in gene expression result in different polarization states.

Here, we used a high-throughput mRNA sequencing for gene expression analysis using the *cdc10-129* mutant to identify cell-cycle dependent factors that regulate the transition from monopolar to bipolar growth. We compared the gene expression profiles of cells in the G1/S and G2 phases using the DESeq2 R package. Gene enrichment analyses and protein-protein interaction network indicate that cells in the G1/S phase of the cell cycle show upregulation of specific stress response pathway genes involved in G1-arrest. On the other hand, the G2 phase shows upregulation of genes involved in plasma membrane modulation and cell-wall biosynthesis. We further evaluated the cell-cycle-dependent expression patterns of these genes via qPCR. In agreement with our gene expression analyses, we find that the stress kinase *spk1*, upregulated in G1, prevents precocious bipolar growth, while the genes *cdc15* and *hob3,* upregulated in G2, prevent monopolar growth. Our data indicate that nutritional stress response/G1-arrest pathways are upregulated in the G1/S phase. In the absence of nutritional stress, these pathways are downregulated to not only promote cell cycle progression but also promote bipolar growth. In the G2 phase, the pathways that promote extensile growth, including plasma membrane and cell wall organization, are upregulated. Our data indicate that cell-cycle phase-dependent differential gene expression promotes the transition from monopolar to bipolar growth.

## Results

### Cell-cycle-dependent RNA isolation in *cdc10-129* mutants for gene expression analysis

Cell-cycle progression of *cdc10-129* mutant cells was performed using canonical cell-cycle arrest and release experiments. *cdc10-129* mutant cells growing in early-log phase at 25°C was arrested in the G1/S phase at 36°C for 4 hours. The cells were then released to 25°C and monitored for cell-cycle progression using DAPI and calcofluor staining. We observed that cells arrested at 36°C are monopolar (**Supplementary Figure S1A**). When released to 25°C, more than 90% of the cells are bipolar after 2 hours and 30 minutes (**Supplementary Figure S1B**), and bi-nucleate after 3 hours, suggesting entry into mitosis. This indicates that 2 hours and 30 minutes after release, the cells are synchronized in G2 phase.

The mRNA experimental design workflow for cell synchronization using *cdc10-129* mutants is shown in **Supplementary Figure S2A**. Early-log phase cells growing at 25°C (asynchronous), arrested at 36°C (G1/S), and post-release at 25°C (G2) for 2 hours and 30 minutes were collected for RNA isolation. Similar conditional controls with wild-type cells were also collected for RNA extraction and high-throughput sequencing. The isolated RNA was used for library construction, quality control, and sequencing as per the workflow shown in **Supplementary Figure S3A and S3B**. We next analyzed the quality of the RNA sequencing data by estimating the error rate distribution, GC content distribution, and data filtering. In our quality control analysis, we found that the single-base error rate for all 18 samples was lower than 1% (**Supplementary Table S5**). Analysis of GC content distribution showed equal and stable G-C and A-T content throughout the sequencing process for all 18 samples (**Supplementary Table S6**).

### Differential expression of gene analysis shows cell-cycle-dependent differences in gene regulation

The gene expression data were statistically analyzed using the DESeq2 R package between the different groups. Variability between biological replicates was calculated using edgeR to determine the variability within biological replicates. Post statistical analysis, the alignment of the sequence was performed using HISAT2 with the reference genome (**Supplementary Table S7**). Mapped regions are classed as exons, introns, or intergenic regions. The distribution of sequencing reads across all samples in the genomic area was found to have 97-99.5% exons, and the remaining area belonged to either introns or intergenic regions. The gene expression data for all 18 samples from 3 independent samples are summarized in **Supplementary Table S8**. Distribution of gene expression levels under different conditions and fragments per kilobase per million mapped fragments (FPKM) among different samples is represented as boxplots shown in **Supplementary Figure S4B**. For biological replicates, the final FPKM is represented as the mean value.

We used Pearson correlation coefficient to calculate the differential expression between the samples. A correlation coefficient closer to 1 indicates high similarity within the samples. As per ENCODE [35] (ENCODE Project Consortium, 2004), the square of the Pearson correlation coefficient should be greater than 0.92, and the R^2^ value should be greater than 0.8. We calculated the inter-sample correlation coefficient R^2^ value within the range of 0.8 to 1 (**Supplementary Figure S5A**).

To evaluate the intergroup differences and intragroup sample duplication, we utilized the principal component analysis (PCA) method. We performed PCA analysis on the gene expression value (FPKM) of all samples (**Figure 1A**). We find a distinct clustering pattern between *cdc10-129* cells growing in G1/S and G2 phase of the cell cycle, with principal components 1 and 2 (PC1 and PC2) accounting for 32.56% and 22.82% of the total variance, respectively. While the G1/S and G2 phase cells showed distinct clusters, one biological replicate in each group did not cluster as tightly, indicating some degree of biological variability within these groups. All other conditional controls from *wild-type* cells and the early log-phase *cdc10-129* cells that were asynchronous in different cell-cycle stages were widely dispersed. Together, these data suggest higher gene expression similarities within given synchronized *cdc10-129* samples and a wide variety of differential gene expression in the asynchronous control samples.

**Figure 1:**
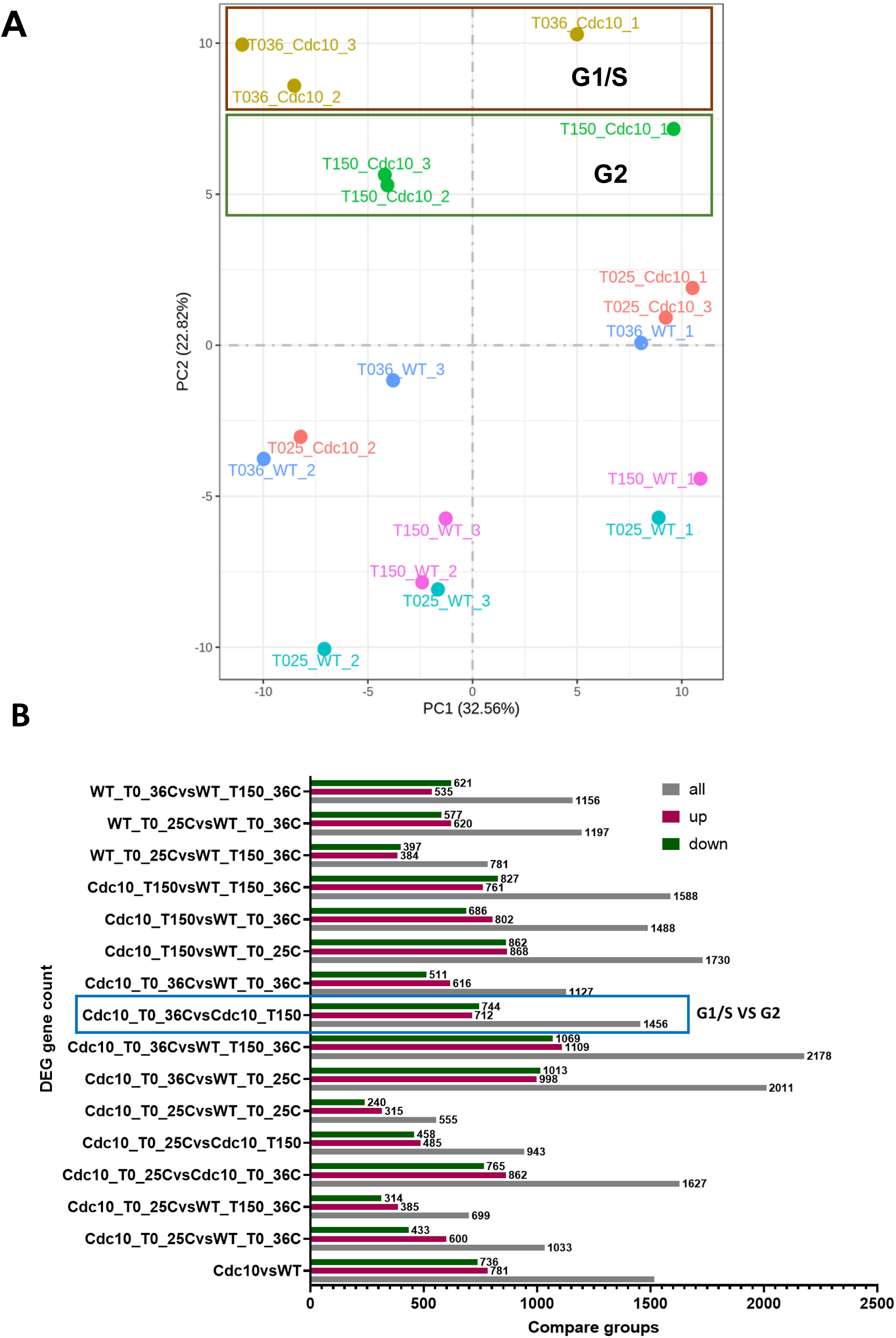
Quality control of high-throughput mRNA sequencing data. A) Principal component analysis result representing intergroup differences and intragroup sample duplication. The orange and green rectangle highlights the G1/S and G2 phases, respectively, of the *cdc10-129* mutant cells. B) Histogram for the number of differential genes for each comparison combination. Pink and green represent the differential genes for up-regulation and down-regulation, respectively, and the numbers on the columns indicate the differential genes.

For the differential expression of gene analysis, we calculated the statistics and threshold for the number of differential genes between different groups (**Figure 1B**). We observe that out of the 1456 differentially expressed genes, 712 are up-regulated in G1/S phase, while 744 were up-regulated in the G2 phase.

The differential gene set was generated by combining all the genes in the comparison group that showed differential expression. Cluster analysis was performed on distinct gene sets for more than two groups, whereas genes with comparable expression patterns were grouped. We used mainstream hierarchical clustering to cluster the FPKM values of genes and homogenized the row (Z-score). Genes from 18 different samples, pooled into 6 independent groups, were used to generate the heatmap (**Figure 2**). Genes that have a similar pattern of expression were grouped on the heatmap. Each grid’s color represents the value derived after homogenizing the expression data, typically between −2 and 2. Our clustering data heatmap (**Figure 2**) showed inter-group clustering and also inter-sample clustering. We observe a clear distinction in the gene expression clustering profile for the cells in G1/S phase and the G2 phase cells. The clear demarcation of the clustering profile in these cells indicates that different cell cycle stages show differential gene expression.

**Figure 2:**
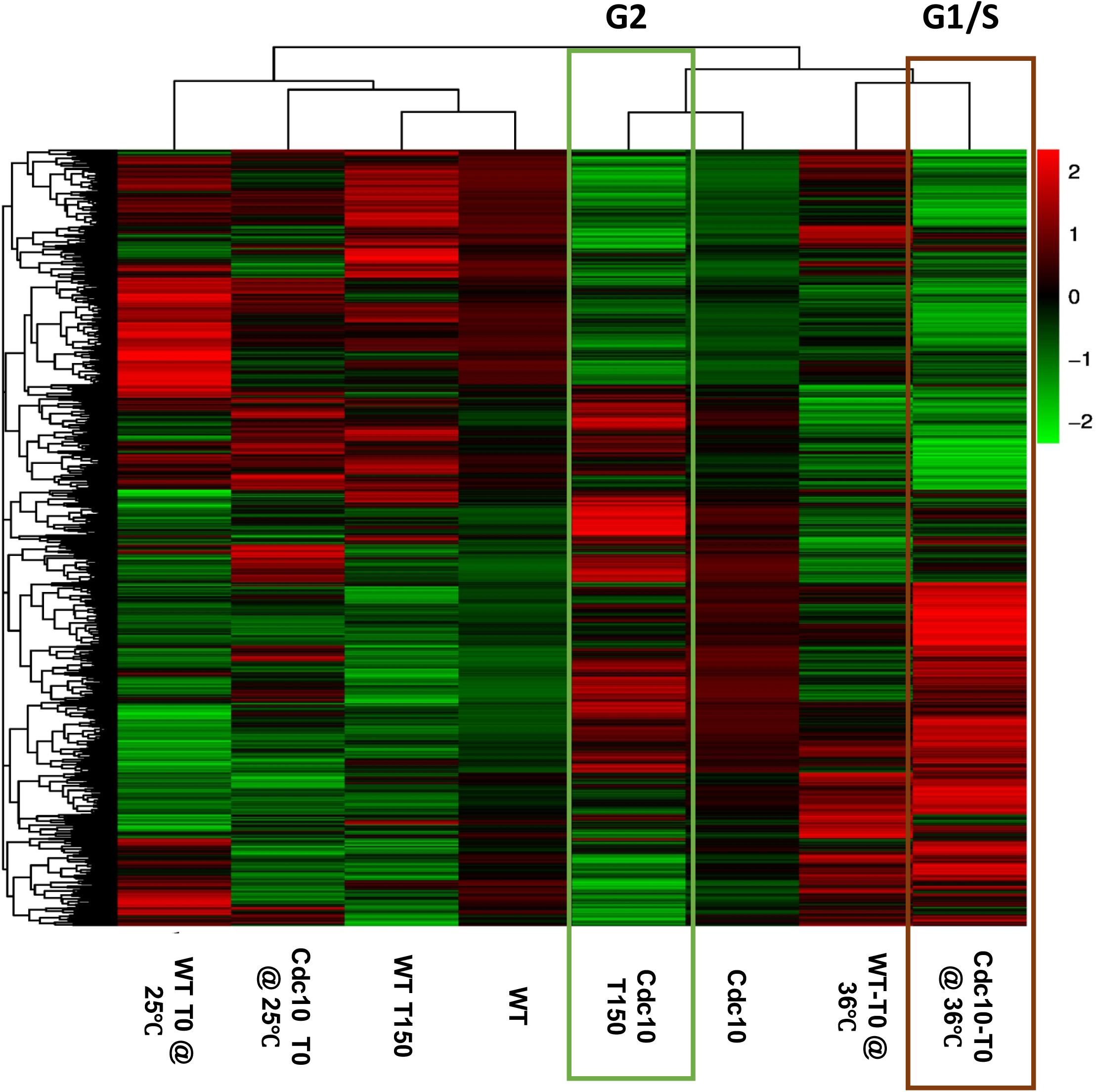
Heat-Map representation of differential gene clustering. The overall results of FPKM cluster analysis using the log2(FPKM+1) value. Red indicates genes with high expression levels, and green indicates genes with low expression levels. The red and green boxes highlight a heatmap representing differential gene expression in G1/S and G2 phases of the *cdc10-129* mutant cells.

The co-expression Venn diagram shows that the overlapping region between different *cdc10-129* samples contains a total of 4874 genes (**Supplementary Figure S5B**). The uniquely identified genes for the three different *cdc10-129* samples, namely asynchronous *cdc10-129* at 25°C, G1/S phase-arrested at 36°C, and G2 phase post-release at 25°C, are reported as 13, 65, and 35 genes, respectively. 87 genes are co-expressed between the G1/S phase cells and the G2 phase cells.

For different experimental conditions, we selected appropriate software for differential gene expression analysis (**Supplementary Table S3**). We screened genes using a threshold of log2[Foldchange]≧1 and adjusted p-value, padj≦0.05. For our analysis, we were only interested in the expression of genes that were differentially expressed between the G1/S phase and the G2 phase of the cell cycle. Therefore, we filtered the enrichment results from the raw differential analysis table to identify only the genes of interest (**Supplementary Table S9**). The goal of this investigation was to identify gene expression patterns that drive the transition from monopolar to bipolar growth in a cell-cycle-dependent manner. Hence, in our analysis, we selected genes that have been shown to play a role in cell polarity. To substantiate our findings, when the number of differential genes discovered was too large, we screened using a stringent threshold criterion and lowered the adjusted p-value (FDR) to ≦0.05.

We identified several genes differentially expressed between the G1/S and G2 phase of the cell cycle. These genes, along with their log2fold change and significant pathway, are reported in **Table 1**. Genes such as *rgs1*, *spk1*, *hsp90*, and *ste11* showed upregulation in the G1/S phase. In the G2 phase cells, genes such as *gas1*, *gas5*, *lat1*, *pda1*, *pkd2*, *hob3*, and *cdc15* were upregulated. Our data show that the majority of genes upregulated in the G1/S phase are involved in regulating the MAPK pheromone response pathway, protein processing in the endoplasmic reticulum, response to pheromone, and conjugation. Genes upregulated in the G2 phase cells are involved in cell-wall organization, plasma membrane maintenance, and glucose metabolism.

**Table 1:**
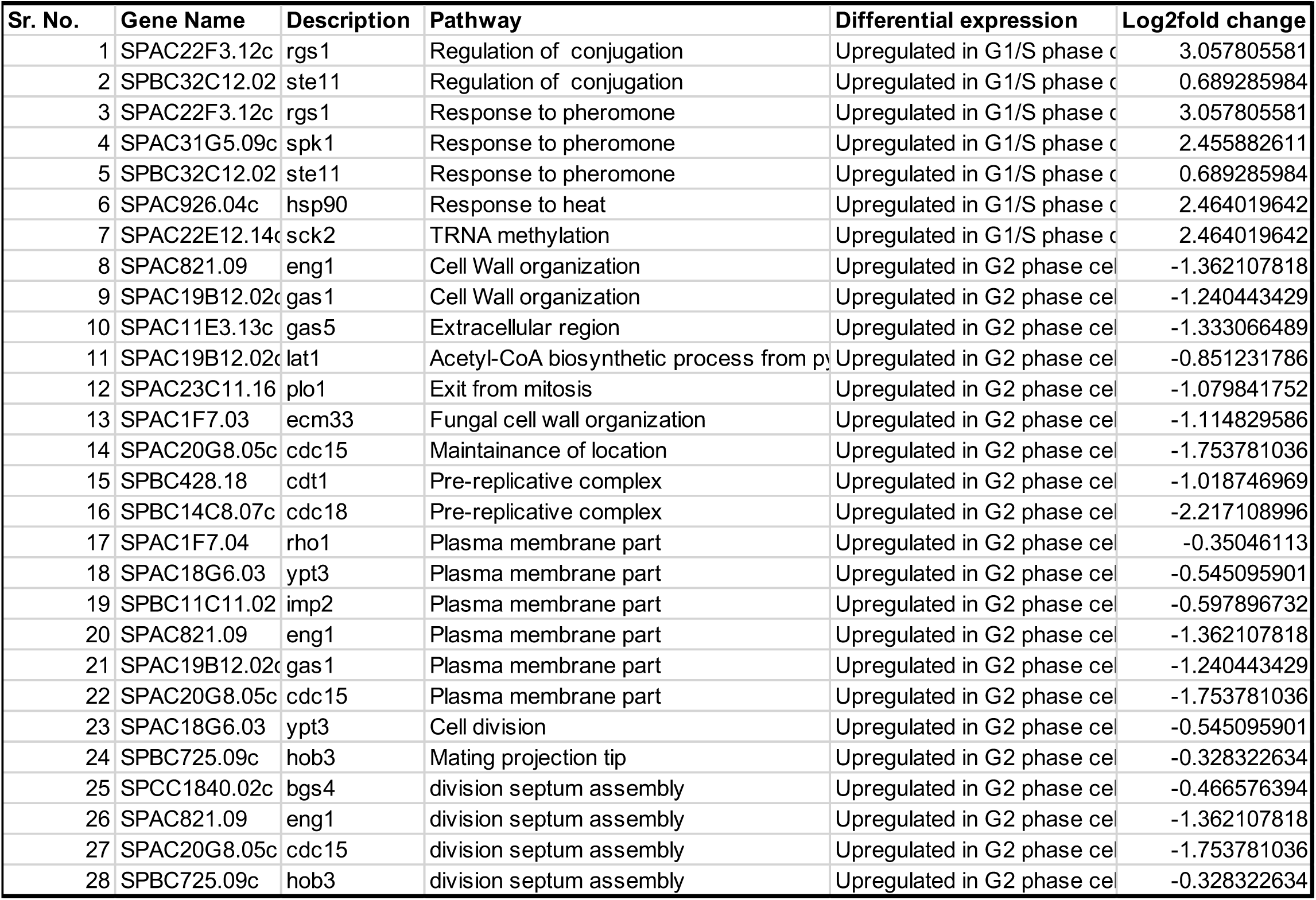
Differential expression of gene between G1/S VS G2 cells in essential pathway based on Log2fold change, P value < 0.05. The values for the log2fold change are indicated for all the genes. “+” indicates up-regulated and “-” indicates down-regulated genes.

### GO and KEGG enrichment analysis of differentially expressed genes identifies pathways for polarity control

Next, we performed Gene Ontology or GO enrichment analysis (http://www.geneontology.org/) to present gene properties of the cellular component, molecular function, and biological process across all species. GO terms with padj-value≦0.05 are considered significantly enriched. In our GO enrichment analysis, the 30 most significant terms were selected for each comparison group. The data for GO terms for cells growing in the G1/S phase and G2 phase are shown in **Figure 3**. We have also reported in **Supplementary Figures S6 and S7**, the GO terms upregulated for G1/S and G2 phase cells. We observe that most of the GO terms associated with differentially expressed genes between G1/S and G2 phase cells regulate various biosynthetic and metabolic pathways, including cell-wall organization and plasma membrane maintenance. The pathways associated with differentially expressed genes in G1/S phase cells were involved in regulating protein folding, rRNA processing, cellular response to pheromone, 90s ribosome, unfolded protein binding, and heat-shock protein binding. While in the G2 phase, we observed genes upregulated that control oxalic acid metabolism, organic acid metabolism, fungal cell wall organization, organelle and endoplasmic reticulum sub compartment, cofactor and coenzyme binding.

**Figure 3:**
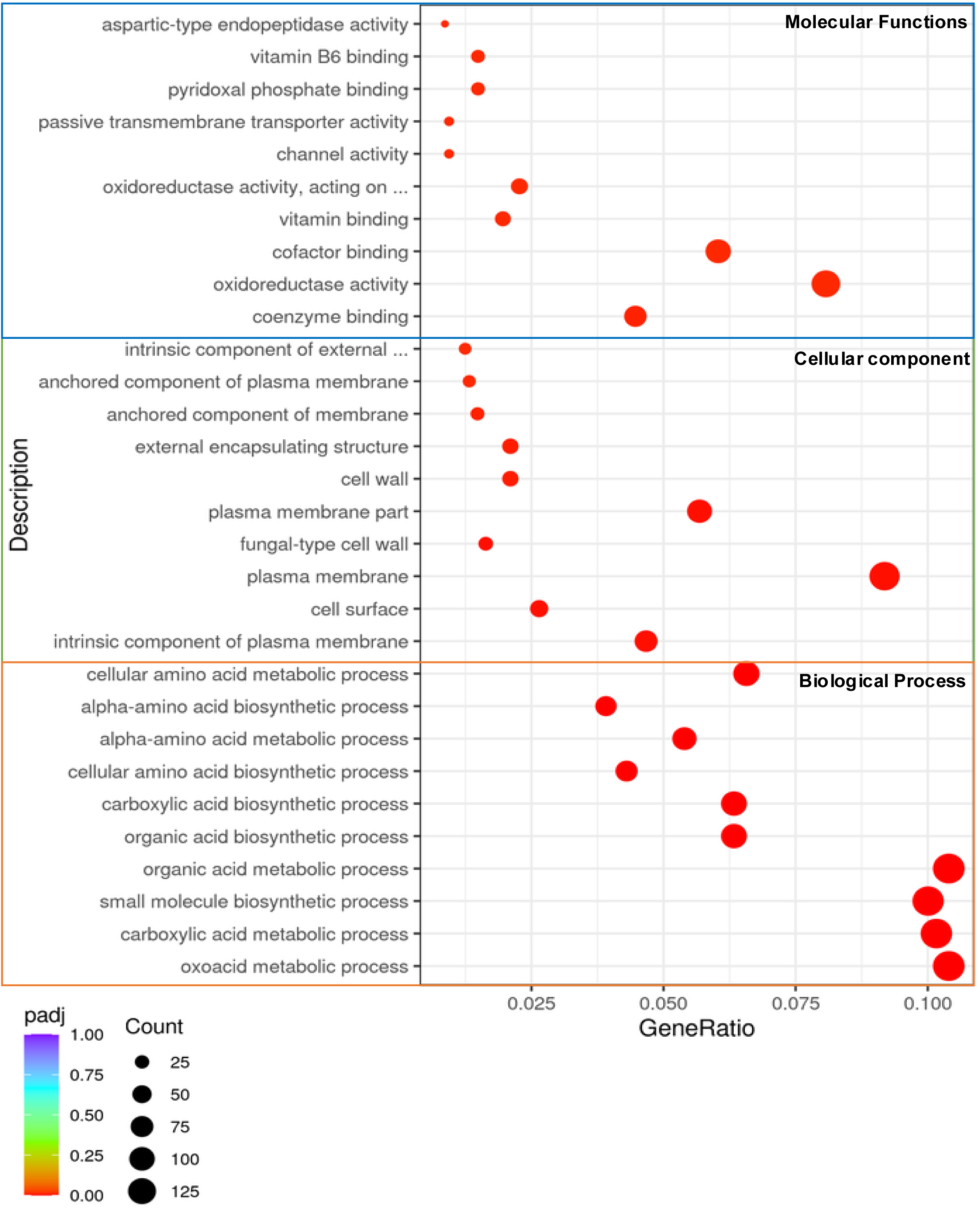
GO enrichment analysis: GO Enrichment Analysis Scatter Plot with the 30 most significant GO terms in G1/S vs G2 population. The abscissa is the ratio of the number of differential genes linked with the GO Term to the total number of differential genes, and the ordinate is the GO Term. The size of the point represents the number of genes annotated to a specific GO Term, and the color from red to purple represents the significance level of the enrichment. The three different categories of biological process, cellular component, and molecular function are represented with orange, green, and blue outlines.

The KEGG enrichment analysis for cells growing in G1/S and G2 phase of the cell-cycle displays a scatter plot of up to 20 of the most significant pathways, differentially regulated in G1/S phase cells and G2 phase cells (**Figure 4**). 20 upregulated pathways for genes associated with G1/S phase and G2 phase of the cell cycle are reported in **Supplementary Figures S8 and S9 and Supplementary Table S10 and S11**. Some of the important pathways with upregulated genes for G1/S phase include protein processing in the endoplasmic reticulum, biosynthesis of cofactors, vitamin B6 metabolism, and RNA degradation. Upregulated genes associated with the G2 phase of the cell cycle include the TCA cycle and various metabolic pathways. These data suggest that cells growing in the G2 phase regulate the metabolism of glucose and various amino acids to prepare the cell to undergo bipolar growth in the presence of available nutrients.

**Figure 4:**
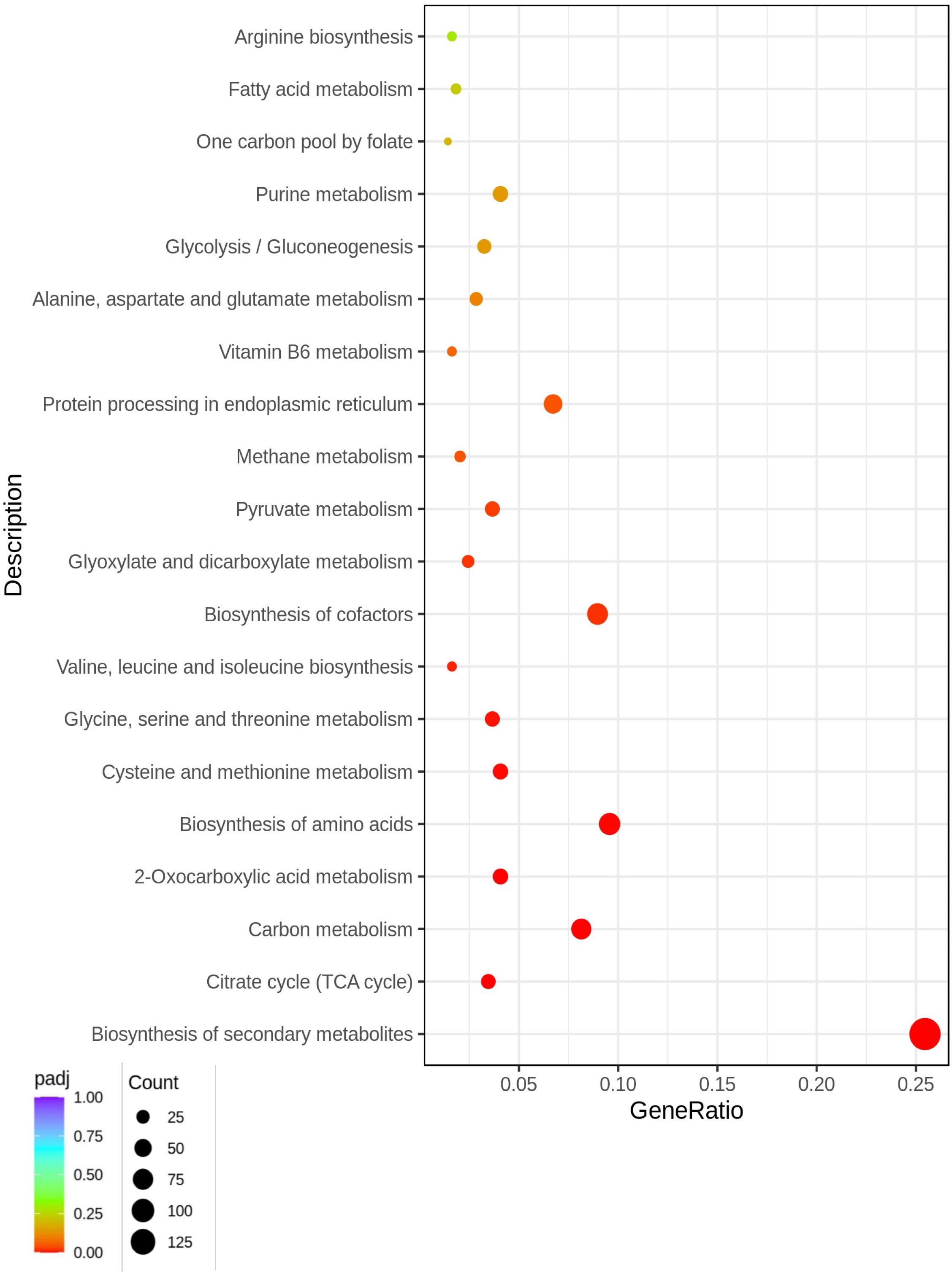
KEGG Pathway analysis: KEGG enrichment scatter plot with 20 most significant pathways showing differential expression in G1/S and G2 phase cells. The abscissa in the graph is the ratio of the number of differential genes on the KEGG pathway to the total number of differential genes, and the ordinate is the KEGG pathway. The size of the point represents the number of genes annotated to a specific GO Term, and the color from red to purple represents the significance level of the enrichment.

GSEA (Gene Set Enrichment Analysis)[36] analysis was performed for all 16 comparison groups. We have reported upregulated genes in G1/S phase and G2 phase cells in **Supplementary Table S12**. Some of the genes that are differentially regulated in these cell-cycle stages include *hsp90, spk1, rgs1, cdc15, and gas1*. Most of these genes are involved in regulating the pheromone response pathway, metabolic pathway, and pathways controlling the cell wall organization and plasma membrane maintenance. The data obtained via GSEA ranking agrees with our enrichment analysis in GO and KEGG pathway analysis.

### Protein-protein interaction (PPI) analysis of differentially expressed genes suggests clustering of polarity pathways

Protein interaction patterns between differentially expressed genes are identified through the analysis of the STRING protein interaction database [37]. Using the STRING database, we constructed networks that visualize how proteins from differentially expressed genes interact with each other. As observed in **Figures 5 and 6**, we used upregulated genes in G1/S and G2 phases of the cell cycle to provide a network with molecular species represented as nodes and intermolecular interactions represented as edges. Our data for PPI analysis show that the genes upregulated in G1/S and G2 phase have interacting partners in either upstream or downstream pathways controlling cell polarization. These genes likely work in concert to globally regulate cell polarization in a cell-cycle-dependent manner. Upon further analysis of PPI for G1/S and G2 phase cells, we identified the biological processes, cellular components, and molecular functions similar to those observed in our GO analysis (**Supplementary Figure S10-S13**). We identified upregulation of the *spk1-byr2-ste11* hub in G1/S phase cells (**Supplementary Figure S10**). These genes are involved in pheromone response and G1-arrest due to nutritional starvation [38, 39]. The biological processes, molecular functions and cellular components upregulated in G1/S are listed in **Supplementary Figure S11**. In the G2 phase cells we identified the *cdc15*-*hob3*-*rho1*-*bgs1* hub that is involved in regulating pathways responsible for cell-wall organization **Supplementary Figure S12**. The biological processes, molecular functions and cellular components upregulated in G2 identified pathways in positive regulation of cellular component biogenesis, plasma membrane maintenance, fatty acid metabolism, and TCA cycle regulation (**Supplementary Figure S13**) [22, 40–43]. These findings indicate upregulation of pathways that promote cell growth and expansion in G2.

**Figure 5:**
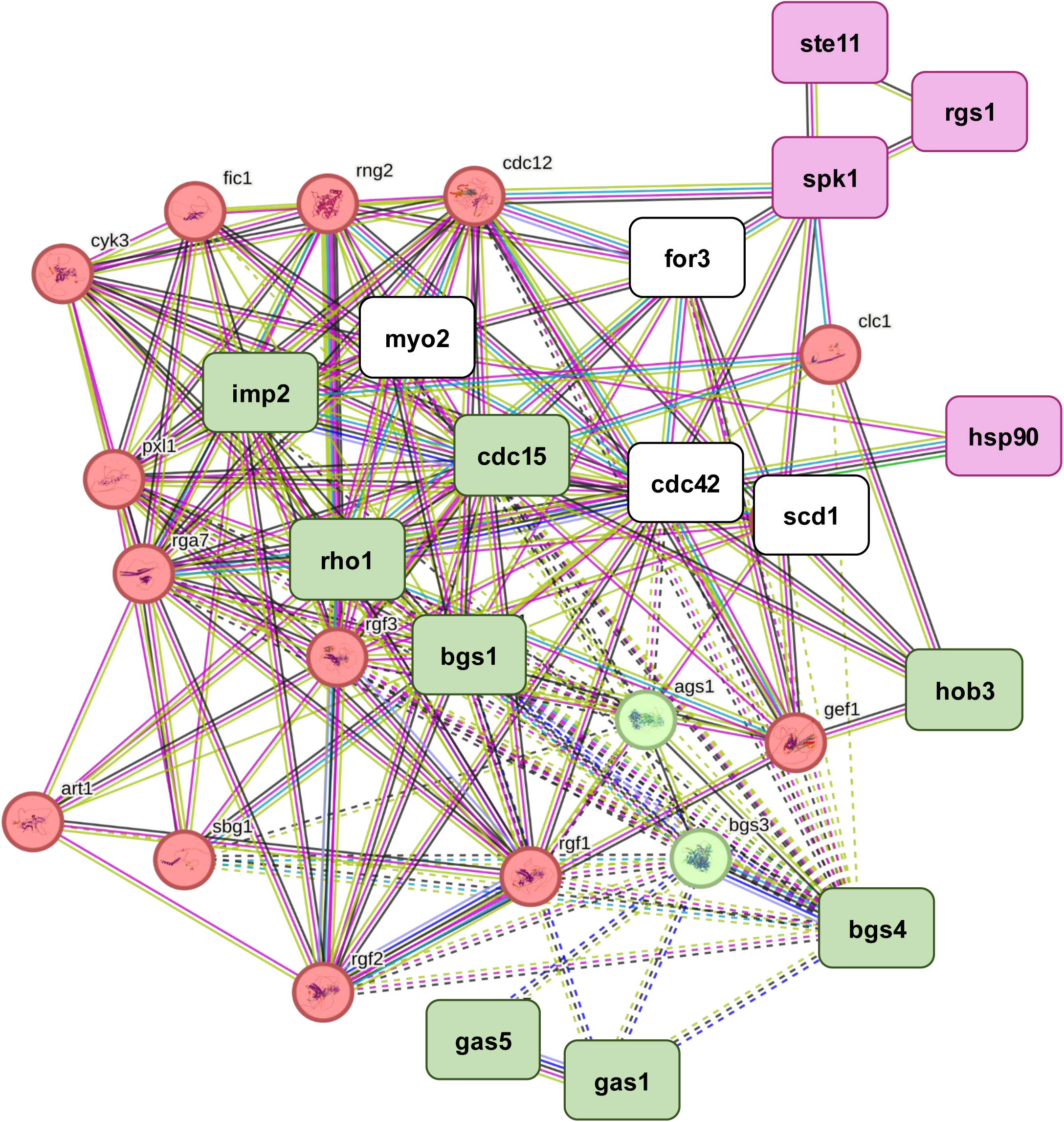
Protein-protein interaction analysis for upregulated and down-regulated genes in the G1/S phase cells. Genes highlighted in green were down-regulated, whereas those in pink were upregulated in our G1/S phase cells when compared to G2 phase cells. There are a total of 31 nodes, including 189 edges, in the figure representing both the down-regulated and up-regulated genes in the G1/S phase cells. The average node degree was estimated to be 12.2, whereas the average local clustering coefficient is 0.755. PPI enrichment value was estimated to be < 1.0e-16. Three basic clusters appear in the PPI analysis using k-means clustering. Red outline around the gene indicates genes involved in cytokinetic process, cell-cortex region, contractile ring formation, and small GTPase-mediated signal transduction. Green outline around the gene indicates genes involved in fungal-type cell-wall polysaccharide synthesis, glycosyl transferase, primary cell septum biogenesis, and UDP glycosyl transferase activity. Blue outline around the gene indicates genes involved in fungal-type cell wall (1->3)-beta-D-glucan biosynthesis.

**Figure 6:**
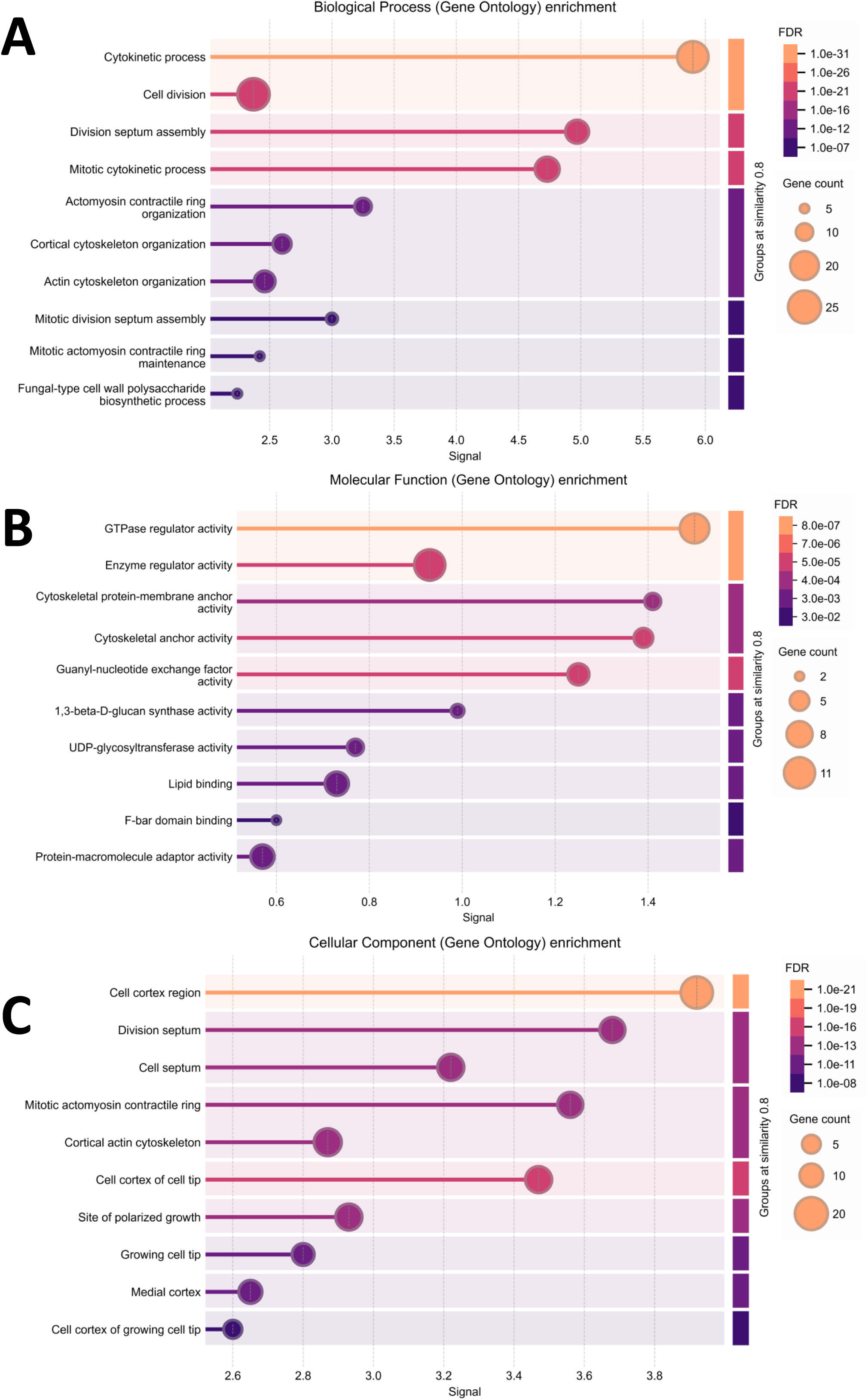
Protein-protein interaction analysis using string for down-regulated and up-regulated genes in the G1/S phase *cdc10-129* cells. Top 10 Gene Ontology enrichment pathways identified by protein-protein interaction analysis in the following categories: A) Biological Process, B) Molecular Functions, C) Cellular Component.

### Cell-cycle-dependent gene expression patterns correspond to cell polarization patterns

Our bioinformatics data identified two Bin-Amphiphysin-Rvs (BAR) domain-containing genes, the F-BAR gene *cdc15* and the N-BAR gene *hob3*. The F-BAR *cdc15* is required for cytokinesis and is also a component of the endocytic complex [22]. Hypomorphic mutants of *cdc15* are monopolar, while hypermorphic mutants are precociously bipolar [22]. Cdc15 promotes bipolarity via the recruitment of the Cdc42 GEF Gef1 to the growing ends [22]. The *hob3* gene is also involved in Gef1-mediated Cdc42 activation [44, 45]. Cells lacking *hob3* mislocalize Gef1 and show decreased Cdc42 activation. Gef1 helps establish Cdc42 positive feedback activation and is required for new-end-take-off leading to bipolar growth [20, 22, 44]. Thus, the upregulation of *cdc15* and *hob3* in the G2 phase of the cell cycle agrees with their role in promoting bipolar growth.

Our RNA seq data analysis also showed that the MAPK pheromone response pathway is upregulated in the G1/S phase. We identified the MAP kinase Spk1 as one of these upregulated genes. The gene *spk1*, a nutritional-stress-related MAP kinase, regulates response to pheromone, conjugation, and mating pathways [38, 39, 46, 47]. Spk1 kinase also promotes G1 to G0 transition in response to nitrogen starvation [39]. Thus, it is expected that this pathway, including Spk1, is upregulated in G1 phase cells. However, it is unclear if this pathway is involved in the transition from monopolar to bipolar growth. We asked if upregulation of *spk1* in G1 prevents monopolar to bipolar transition. Indeed, we find that in *spk1*Δ mutant cells at 25°C, about 80% of the cells were bipolar compared to *spk1+* cells with 51% bipolar cells (**Figure 7**). This indicates that in the absence of the *spk1* gene, cells initiate bipolar growth prematurely. This suggests that the high *spk1* levels in G1/S phase cells prevent the transition to bipolar growth.

**Figure 7:**
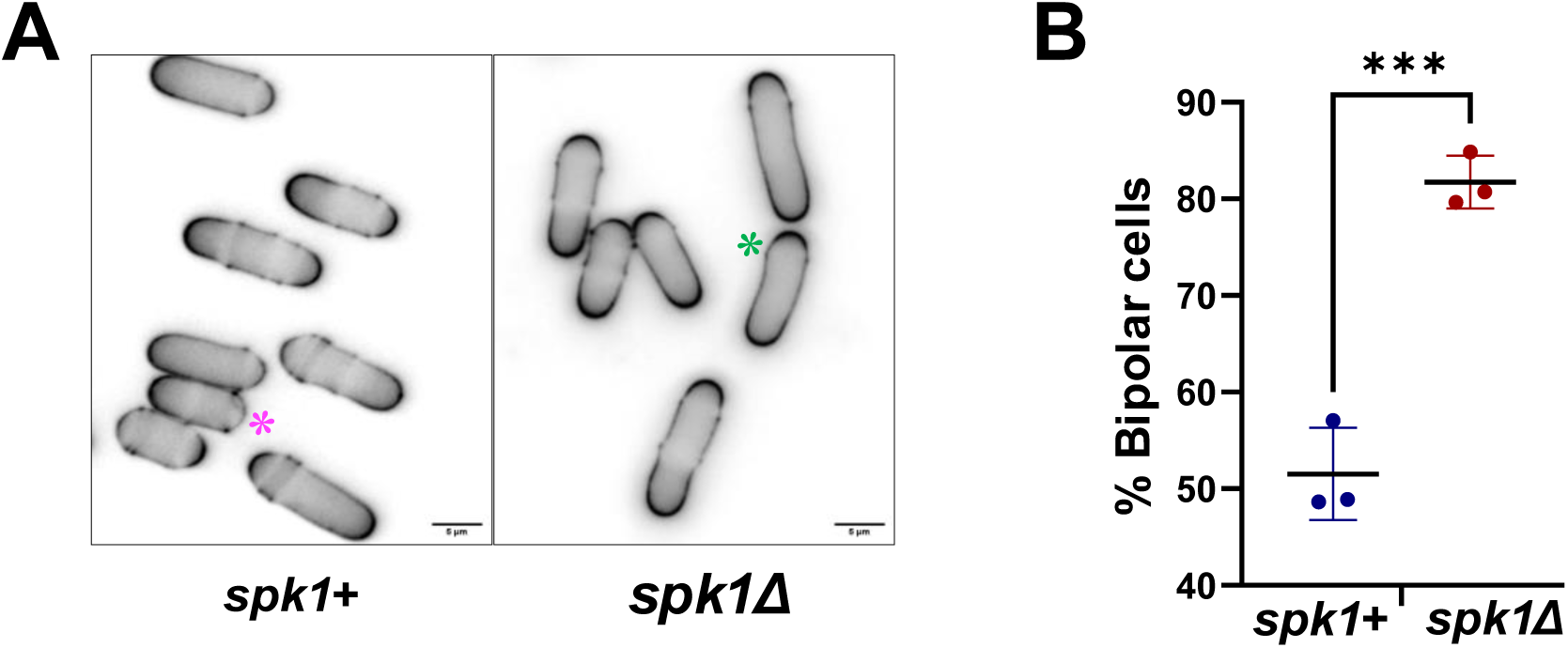
The MAP kinase *spk1* prevents precocious bipolar growth. A) *spk1+* and *spk1*Δ cells at 25℃. B) Quantitative analysis of percentage bipolar cells in *spk1+* cells and *spk1*Δ at 25°C, magenta asterisk represents the monopolar cells, green asterisk represents bipolar cells in *spk1+* cells and *spk1*Δ. Statistical analysis, unpaired t-test. *** P<= 0.0007.

Based on our comprehensive analysis of the gene expression data and our observation with *spk1*Δ cells, we hypothesize that the MAPK pheromone and nutritional stress response pathway is closely linked to the transition from monopolar to bipolar growth. In the G1 phase, the stress pathway is upregulated by default, leading to monopolar growth. In the absence of nutritional stress, the G1 checkpoint is completed, and cells enter the S phase and subsequently the G2 phase. The downregulation of the stress pathway in G2 allows the cells to transition to bipolar growth.

### Validating cell-cycle-dependent gene expression profiles of polarity pathways

To obtain cell-cycle-specific mRNA, we used the *cdc10-129* temperature-sensitive mutant. In our experimental design, the mutant at high temperature (36°C) is arrested in G1/S and released to G1 when grown under permissive conditions (25°C). It is possible that the difference in the temperature conditions impacted gene expression in the *cdc10-129* mutants. However, KEGG analysis of our data in wild-type cells grown at 25°C compared to 36°C displayed very different pathways compared to our data from the G1/S and G2 phase cells (**Supplementary figure S14**). We identified close to 95 statistically significant temperature-responsive transcripts (p < 0.05). Gene ontology enrichment analysis revealed significant changes in genes associated with ribosomal functionality and SNARE-mediated vesicular transport mechanisms. Notably, no statistically significant alterations were observed in transcripts governing MAPK signaling cascades, cell-cycle regulatory networks, or cell polarity determinants in wild-type specimens.

To further validate our gene expression data, we performed qPCR for *spk1*, *cdc15,* and *hob3* with reference to the β-actin gene *act1* in *wild-type* cells and synchronized *cdc10-129* cells (G1/2 at 36°C and G2 at 25°C) [48]. The *act1* gene demonstrates remarkable transcriptional stability across diverse physiological conditions and hence was selected as the qPCR reference gene. Our high-throughput RNA sequencing analysis revealed minimal expression variability (coefficient of variation <0.1) across experimental conditions, regardless of temperature and cell-cycle phase. Statistical analysis of normalized read counts demonstrated no significant differential expression (p>0.05, FDR-adjusted) of *act1* under any tested condition. This robust transcriptional stability, coupled with its abundant expression levels (FPKM >500), establishes *act1* as an optimal internal control for relative quantification in the qPCR analyses. Using qPCR, we show that in the wild-type cells, irrespective of the temperature, *cdc15*, *hob3,* and *spk1* levels do not change. In contrast, *cdc10-129* mutants in G2 phase show upregulation of *cdc15* and *hob3* (**Figure 8A and 8B**). Similarly, during the G1/S phase in the *cdc10-129* mutant *spk1* gene was upregulated (**Figure 8C**). Together, these results further indicate that the differential gene expression data in G1/S and G2 phase cells is not due to the effects of the different temperatures.

**Figure 8:**
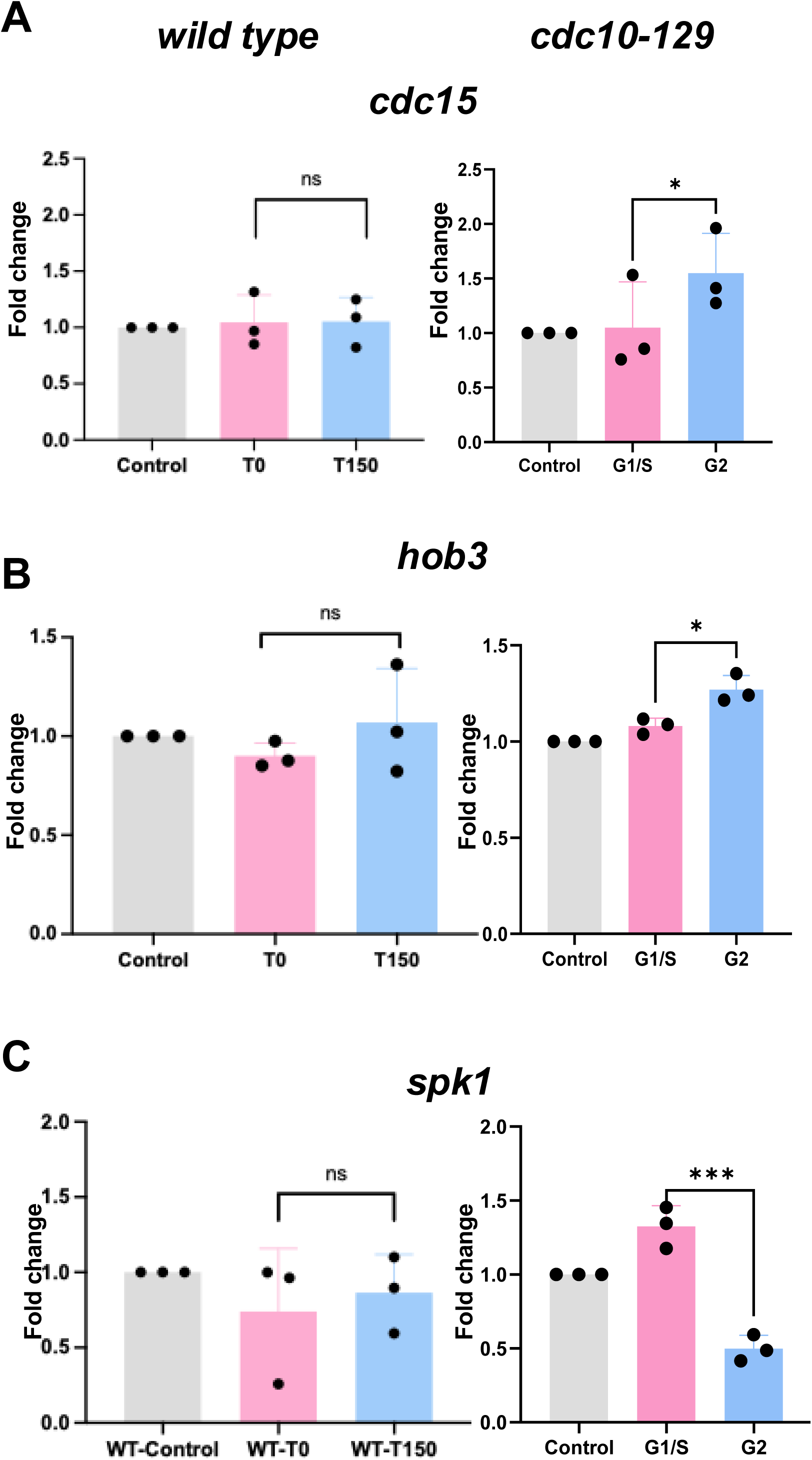
qPCR expression profile for *cdc15, hob3,* and *spk1* in *wild-type and cdc10-129* mutant cells. Fold change in mRNA expression in *wild-type and* cells in different cell-cycle stages for *cdc15* (A), *hob3* (B), and *spk1* (C). Grey bar represents control asynchronous cells in *wild-type* and *cdc10-129* mutant cells at 25°C. The pink bar represents G1/S phase cells in *cdc10-129* mutant cells and asynchronized *wild-type* cells growing at 36℃. The light blue bar represents G2 phase cells in *cdc10-129* mutant cells, as well as asynchronous *wild-type* cells at 2 hours and 3 hours after shift to 25°C. The error bar represents the standard deviation, n=3 replicate experiments. Statistical analysis between the two conditions was performed by paired Student’s t-test, n.s. is not statistically significant, * P<0.05, ***P< 0.0007.

## Discussion

The initial step in cell polarization follows cues generated post-cell division, leading to spatiotemporal organization of polarity proteins for the regulation of the cytoskeleton and cell growth [2, 8, 10, 49–54]. In fission yeast, spatiotemporal regulation of Cdc42 drives polarized growth. Despite the small subset of proteins regulating Cdc42 self-organization at the growth site [18, 20], the underlying principles of cell-cycle-dependent polarized growth remain unclear.

In the current work, we explore the major pathways controlling the transition from monopolar to bipolar growth in a cell-cycle-dependent manner. The transition to bipolar growth only occurs when the cells enter the G2 phase of the cell cycle [55]. To identify the factors that enable this transition, we investigated the gene expression pattern of cell-cycle-arrested G1/S and G2 phase cells using the *cdc10-129* mutant. Our gene quantification analysis showed intragroup clustering in synchronized *cdc10-129* cells in G1/S and in G2 phase. However, there was significant intergroup variability in expression patterns between G1/S phase cells and G2 phase cells. Gene clustering heatmap shows differentially regulated gene expression for G1/S and G2 phase cells. We found that between these specific groups, a total of 712 genes were upregulated in G1/S phase cells, while 744 genes were upregulated in G2 phase cells. Our analysis for the Log2fold expression change showed that genes such as *spk1*, *rgs1*, *hsp90*, *lat1*, *pda1*, *hob3*, *cdc15*, *rho1*, and *pkd2* were differentially expressed between G1/S phase and G2 phase of the cell cycle. Using GO analysis, we categorized the genes into three distinct groups of Biological Process (BP), Molecular Functions (MF), and Cellular Components (CC). Our GO enrichment analysis and KEGG pathway analysis showed that genes upregulated in G1/S phase were involved in controlling protein folding, 90s ribosomes, and rRNA methyltransferase activity. Genes upregulated in G2 phase cells were involved in controlling various metabolic pathways, the TCA cycle, plasma membrane maintenance, cell-wall organization, and transmembrane transporter activity. GSEA analysis was performed to validate whether the prior gene set was significantly different between the two biological states. The GSEA data agreed with the GO enrichment and KEGG pathway analysis.

Protein-protein interaction analysis was performed for the genes with significant differential expression between the G1/S and G2 phases of the cell cycle. Our data shows that genes like *spk1, rgs1, and hsp90* were upregulated in the G1/S phase of the cell, which is upstream of the polarity complex. These genes are known to function in the stress response pathway, where Spk1 is the MAP kinase for nutritional stress [38, 39, 46, 47, 56]. Interestingly, we did not observe cell-cycle-dependent differential gene expression of the stress response MAP kinase Sty1, which promotes depolarization under stress [57–59]. In G2 phase cells, genes such as *hob3* and *cdc15,* which indirectly control Cdc42 activation via Gef1, were upregulated. Hob3 is an N-BAR domain-containing protein that binds and recruits Gef1 to its site of action [19, 60]. Similarly, Cdc15 is an F-BAR domain-containing protein that recruits Gef1 to the site of action. Lack of *hob3* or hypomorphic *cdc15* results in monopolar cells due to the inability of Gef1-mediated Cdc42 activation in these mutants [22, 60].

The cell-cycle-dependent gene expression patterns of the nutritional stress response pathways are in agreement with the fact that the G1-G2 phase transition in the cell cycle only occurs in the presence of nutritionally rich conditions [55]. Cells growing in the G1/S phase can have two different fates depending upon the environment they are in: either they enter the S phase and eventually the G2 phase in a stress-free environment, or they enter the quiescent G0 phase under nutritional stress. Based on our findings, we hypothesize that the nutritional stress pathways are upregulated by default in the G1 phase of the cell cycle, where the cells are monopolar. In nutritionally rich conditions, stress pathways are downregulated, and the cells enter G2, resulting in bipolar growth. Indeed, we find that in cells lacking *spk1,* the transition to bipolar growth occurs prematurely, indicating that *spk1* expression promotes monopolar growth.

Cell-cycle-dependent regulation of cell polarity is likely mediated by multiple regulatory modules. While our data identifies how gene expression patterns regulate the transition from monopolar to bipolar growth, we do not rule out the possibility of the role of cell-cycle-dependent kinases in this process. However, it should be noted that even during monopolar growth in wild-type cells, the Cdc42 dynamics at the growing ends are not significantly different from those in bipolar cells [18]. The major regulators of Cdc42, such as the GEFs Scd1 and Gef1, the scaffold Scd2, and the negative feedback kinase Pak1, all localize to the monopolar end, resulting in Cdc42 activation at that end. Our data suggests that the transition to bipolar growth requires a global change in growth regulation rather than specific molecular changes in the Cdc42 regulators. Indeed, our gene expression data show upregulation of cellular component biogenesis, cell wall organization and plasma membrane maintenance pathways in the G2 phase. While the Cdc42 pathway signals and activates growth, for the cell to successfully increase in size, it requires sufficient extension of the plasma membrane and cell wall biogenesis. It is possible that upregulation of these pathways in addition to Cdc42 activation is required for the successful transition to bipolar growth. Further investigations will determine the molecular details of how stress response pathways, changes in the plasma membrane profile, and cell wall reorganization regulate polarity.

Our data indicate that the transition from monopolar to bipolar growth is regulated at multiple levels within the cell. It is conceivable that bipolar growth not only requires activation of the Cdc42 pathway at the two growing ends, but that the cell generates sufficient material via anabolic pathways for the two cell ends to grow effectively. Indeed, we observed upregulation of cell wall biogenesis and membrane remodeling pathways in the G2 phase, where bipolarity occurs. The transition from G1 to G2 requires a nutritional stress-free environment. Here, we show that the nutritional stress response pathway that is known to prevent G1-G2 phase transition also prevents the transition to bipolar growth. In the absence of nutritional stress, these pathways are downregulated and cells transition to the G2 phase. Upregulation of anabolic activities in G2 phase cells promote bipolar growth. Further investigations will reveal whether the stress response pathways down regulate anabolic activity thus preventing bipolar growth. In nature, fission yeast cells typically form pseudohyphae where they grow in a monopolar manner, end-to-end, towards the nutrition source [61, 62]. Under laboratory conditions, fission yeast cells growing in rich media display bipolar growth. Our findings indicate how cell polarity is regulated at the systemic level, where growth conditions and cell-cycle stage play a critical role.

## Materials and Methods

### Strains and genetic methods

All pombe strains used in this study are listed in the **Supplementary Table S1**. The strains used in the study were isogenic to the original strain PN567. The cells were grown in YES (yeast extract) media at 25°C unless otherwise specified. Standard yeast genetic manipulation techniques were used for cell isolation, growth, and analysis [63]. Cells were cultured for three consecutive days for eight generations before imaging studies.

### Cell-cycle progression and synchronization analysis for *cdc10-129* mutant cells

The cell-cycle block and release experiment was performed for strains carrying the *cdc10-129* allele and control wild-type cells[48]. Early log-phase cells were shifted from 25°C to 36°C for 4 h, then shifted back to 25°C for the indicated times. Cell synchronization of the *cdc10-129* mutant was observed every 30 minutes post-release at 25°C through calcofluor and DAPI staining. Microscopic imaging was done on a Nikon Eclipse Ti2 microscope with a 60x objective to determine the percentage of cells with septa and the percentage of bi-nucleated cells. The calcofluor and DAPI imaging was done for 3 independent replicates.

### Cell-cycle synchronization, sample collection, and RNA isolation for high-throughput mRNA sequencing

The early log phase *cdc10-129* mutant and control *wild type* cells were grown at 36°C for 4 hours, followed by release at 25°C for 150 minutes. Cells were collected from samples grown at 36°C and at 25°C for 150 minutes for RNA isolation. The asynchronous controls were also collected for RNA isolation. All 6 different cell types collected were then subjected to centrifugation at 3750 rpm for 10 minutes at 4°C. The supernatant was discarded, while the cell pellet was further used for RNA isolation. RNA isolation was done using the RNeasy Mini Kit from Qiagen (Cat No. 74104) with the standard kit protocol.

### Library construction, quality control, and sequencing

The project workflow is summarized in **Supplementary Figure S2B**. Messenger RNA was extracted from total RNA using poly-T oligo-attached magnetic beads. Following fragmentation, the first strand cDNA was synthesized with random hexamer primers, followed by the second strand cDNA synthesis with either dUTP for a directed library or dTTP for a nondirectional library. The non-directional library was prepared following end repair, A-tailing, adaptor ligation, size selection, amplification, and purification as shown in **Supplementary Figure S3A**. The directional library was prepared following end repair, A-tailing, adapter ligation, size selection, USER enzyme digestion, amplification, and purification, as shown in **Supplementary Figure S3B**. The library was quantified using Qubit and real-time PCR, and its size distribution was detected using a bioanalyzer. Quantified libraries were pooled and sequenced on Illumina platforms based on their effective library concentration and data volume. Based on the wide-ranging evaluation parameters for RNA sample quality requirements for library preparation and sequencing, all 18 samples passed the quality control measures for further analysis and are reported in **Supplementary Table S2**. The detailed workflow for mRNA sequencing data alongside standard bioinformatic analysis with a well-annotated reference genome is shown in **Supplementary Figure S15A**.

### Bioinformatic Analysis Pipeline and Data Quality Control

The workflow for mRNA sequencing data of standard bioinformatic analysis with a well-annotated reference genome is shown in **Supplementary Figure S15A**. Furthermore, we did different bioinformatic analyses, as shown in **Supplementary Figure S4A**, to evaluate the differential gene expression.

CASAVA base recognition (Base Calling) was used to transform the original image data file from a high-throughput sequencing platform, Illumina. Raw data were then stored in FASTQ (fq) format files, which comprised read sequences and base quality values. Raw data (raw readings) were initially processed using Perl scripts. The step, as shown in **Supplementary Figure S4B**, produced clean data (clean reads) by removing adaptors, poly-N, and low-quality reads from raw data. We also computed the Q20, Q30, and GC contents. All subsequent analyses were based on clean data of excellent quality. The adaptor sequence used in the current pipeline is summarized in **Supplementary Figure S4C**. The sequencing error rate for each base was calculated by the Phred score, represented by the equation below,

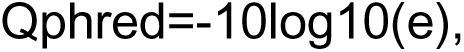

where “e” stands for sequencing error rate, and Qphred represents base quality values obtained from Illumina platforms.

Sequencing reads/raw reads frequently comprise low-quality reads or reads with adapters, reducing the quality of downstream analysis. To circumvent this, the raw reads for our samples were filtered to produce clean reads. The workflow for the sequencing data filtering is shown in **Supplementary Figure S4B**. The adaptor sequence for the same is shown in **Supplementary Figure S4C**.

### Mapping sequencing reads to the reference genome

The PomBase genome website was the source for the reference genome and gene model annotation files [64]. The reference genome index was constructed, and paired-end clean reads were matched to the reference genome using Hisat2 v2.0.5. HISAT2 [65] is a graph-based alignment method that replaced HISAT and TOPHAT2. We selected Hisat2 as the mapping tool since it can generate a database of splice junctions based on the gene model annotation file and generate better mapping results than other non-splice mapping tools.

### Novel transcript predictions

The mapped reads of each sample were assembled by StringTie [66] (v1.3.3b) in a reference-based approach. StringTie uses a novel network flow algorithm as well as an optional de novo assembly step to assemble and quantify full-length transcripts representing multiple splice variants for each gene locus.

### Quantification of gene expression levels

The number of reads mapped to each gene was counted using Feature Counts [67] v1.5.0-p3. The length of the gene and the number of reads mapped to it were then used to compute the FPKM of each gene. Upon quantification of gene expression, statistical analysis for the expression data was performed to screen for the differentially expressed genes in different conditions. The differential analysis was mainly divided into three steps. First, the raw read count was normalized to correct the sequencing depth; followed by applying the statistical model, we calculate the hypothesis test’s probability or p-value. Finally, we utilized multiple hypothesis test corrections to obtain FDR values (false discovery rate) [68]. For different experimental conditions, we selected appropriate software for gene expression differential analysis, as shown in **Supplementary Table S3**.

### Differential expression of gene analysis

Differential expression[68] analysis of two conditions/groups (two biological replicates per condition) was performed using the DESeq2R package [69] (1.20.0). Benjamini and Hochberg’s method for regulating the false discovery rate was used to modify the obtained P-values. Genes with an adjusted P-value ≦0.05 found by DESeq2 were assigned as differentially expressed. Before differential gene expression analysis, for each sequenced library, the read counts were adjusted by the edgeR program package [70] through one scaling factor. EdgeR R package (3.22.5) was used to analyze the differential expression between two conditions. Benjamini & Hochberg method was used to adjust the P-values. An absolute fold change of 2 and a corrected P-value of 0.05 were set as the threshold for significant differential expression of genes.

### GO and KEGG enrichment analysis of differentially expressed genes

Gene Ontology [71] (GO) enrichment analysis of differentially expressed genes was implemented by the clusterProfiler R package, in which gene length bias was corrected. GO terms with a corrected P-value less than 0.05 were considered significantly enriched by differentially expressed genes. We used the clusterProfiler R package to test the statistical enrichment of differential expression genes in KEGG [72] pathways.

### Gene Set Enrichment Analysis

Gene Set Enrichment Analysis (GSEA) [36, 73] was used to determine if a pre-defined Gene set was showing any significant consistent difference between two biological states. All the genes were ranked according to the degree of differential expression in the two samples. The predefined Gene Sets were analyzed to ensure the reliability of enrichment at the top or the bottom of the gene list. Subtle gene expression changes can be analyzed using the GSEA analysis. We used the local version of the GSEA analysis tool http://www.broadinstitute.org/gsea/index.jsp, the GO, KEGG data sets were used for GSEA independently.

### SNP Analysis

GATK [74] (v4.1.1.0) software was used to perform SNP calling. Raw VCF files were filtered with GATK standard filter method together with other parameters (cluster:3; WindowSize:35; QD < 2.0; FS > 30.0; DP < 10. 2.9

### AS analysis

Alternative Splicing is an important mechanism to regulate gene expression and the variability of protein. rMATS[75] (4.1.0) software was used to analyze the AS event.

### Protein-Protein Interaction (PPI) analysis of differentially expressed genes

PPI analysis of differentially expressed genes was performed using the STRING [37] database, with known and predicted Protein-Protein Interactions.

### Cell morphological analysis of the *spk1*Δ mutant

Early log phase cells of control *wild type* and *spk1*Δ cells growing at 25℃ were imaged using calcofluor staining of the cell-wall on the Nikon Eclipse Ti2 microscope at 60X objective. The morphological analysis of percentage monopolar and bipolar cells was calculated using the cell counter feature of ImageJ/Fiji software.

### RNA isolation, cDNA synthesis, and qPCR gene expression analysis of *wild type*, *cdc10-129,* and *cdc25-22* mutant cells

mRNA was extracted from the following samples using Qiagen RNAeasy Mini kit (Cat. No. 74104)

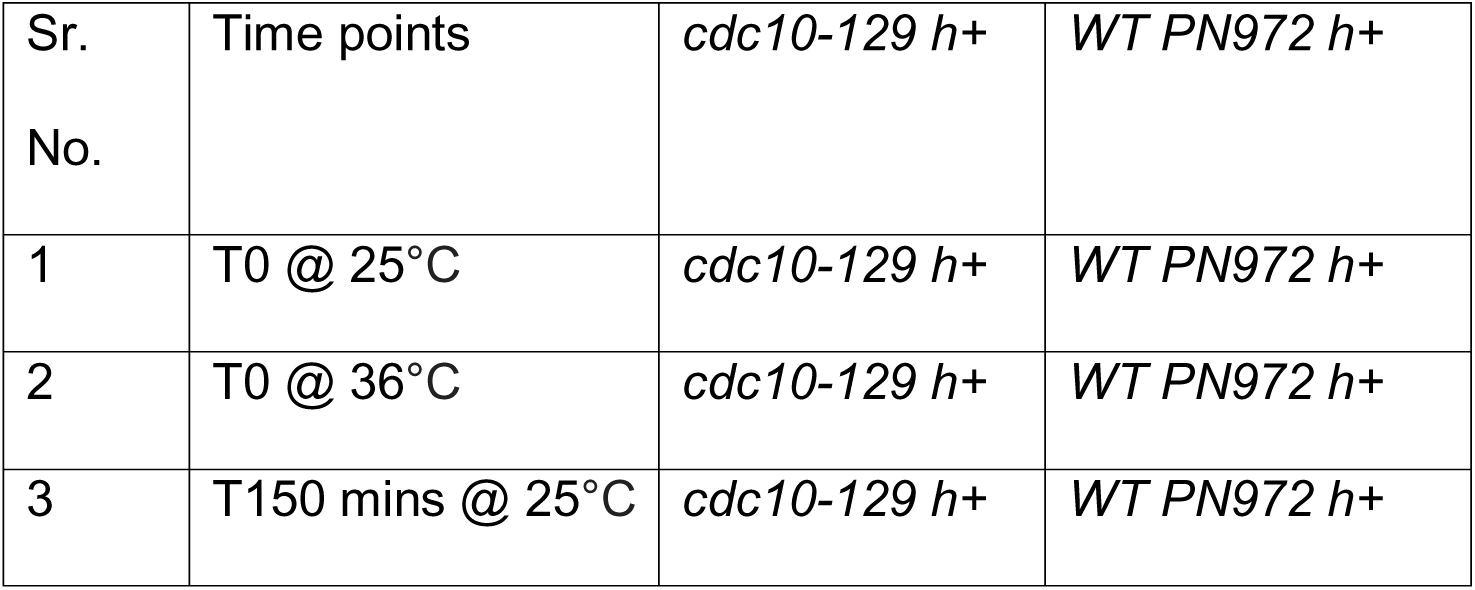

The isolated mRNA was then used to synthesize stable cDNA using NEB LunaScript RT SuperMix Kit (E3010). Real-time quantitative PCR was performed using the NEB Luna Universal One-Step RT-qPCR kit. PCR primers for each of the genes were ordered from IDT (Integrated DNA Technology) and are reported in the **Supplementary Table S4**. The primer efficiency for all the genes, including the housekeeping gene and the gene of interest, was determined by calculating absolute target quantities from an appropriate standard curve, which was derived for each gene from a series of known dilutions. The primer efficiency for each gene was calculated from the standard curve using the equation *y* = *mx* + *c* with an R^2^ value of 0.98

We used dye-based real-time fluorescence of a double-stranded DNA (dsDNA) binding dye, SYBR Green I, to measure DNA amplification after each PCR cycle. Upon fluorescence signal detection over the background fluorescence, a quantification cycle or Ct/Cq values were determined on the QuantStudio Real-Time PCR machine v1.7.2. Ct/Cq values were estimated from three independent biological replicates, which were then used to evaluate the relative target abundance between two or more samples. The expression levels for *hob3*, *cdc15,* and *spk1* were normalized against *act1* housekeeping gene using a robust method[76, 77]. The equation used for log fold change determination is

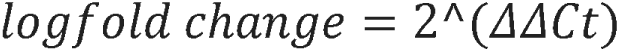

Data were presented as mean ± s.d. Statistical significance of fold change between two conditions was determined using the paired Student’s t test.

## Abbreviations

FPKM: Fragments per kilobase of transcript per million
RPKM: reads per kilobase of transcript per million
qPCR: Quantitative polymerase chain reaction
NETO: New end take off
PCA: Principal component analysis
GSEA: Gene Set Enrichment Analysis
GO: Gene ontology
KEGG: Kyoto encyclopedia of genes and genomes.

## Acknowledgements

The authors would like to thank Boston College for providing the facility to work with pombe systems. We would also like to thank Sophie Martin and Olaf Nielson for providing us with the strains. This work is supported by the National Institutes of Health grant R01GM136847 to M.D.

## Author contributions

Conceptualization: Samridhi Pathak and Maitreyi Das, Methodology: Samridhi Pathak and Maitreyi Das, Analysis: Samridhi Pathak, Validation: Samridhi Pathak and Maitreyi Das, Resources: Maitreyi Das, Funding acquisition: Maitreyi Das, Writing: Samridhi Pathak, Reviewing and editing: Maitreyi Das, Supervision: Maitreyi Das

