## Supplementary Figures S1-S15 for "Differential gene expression drives cell-cycle-dependent transition from monopolar to bipolar growth in fission yeast"

### Supplementary Figure S1

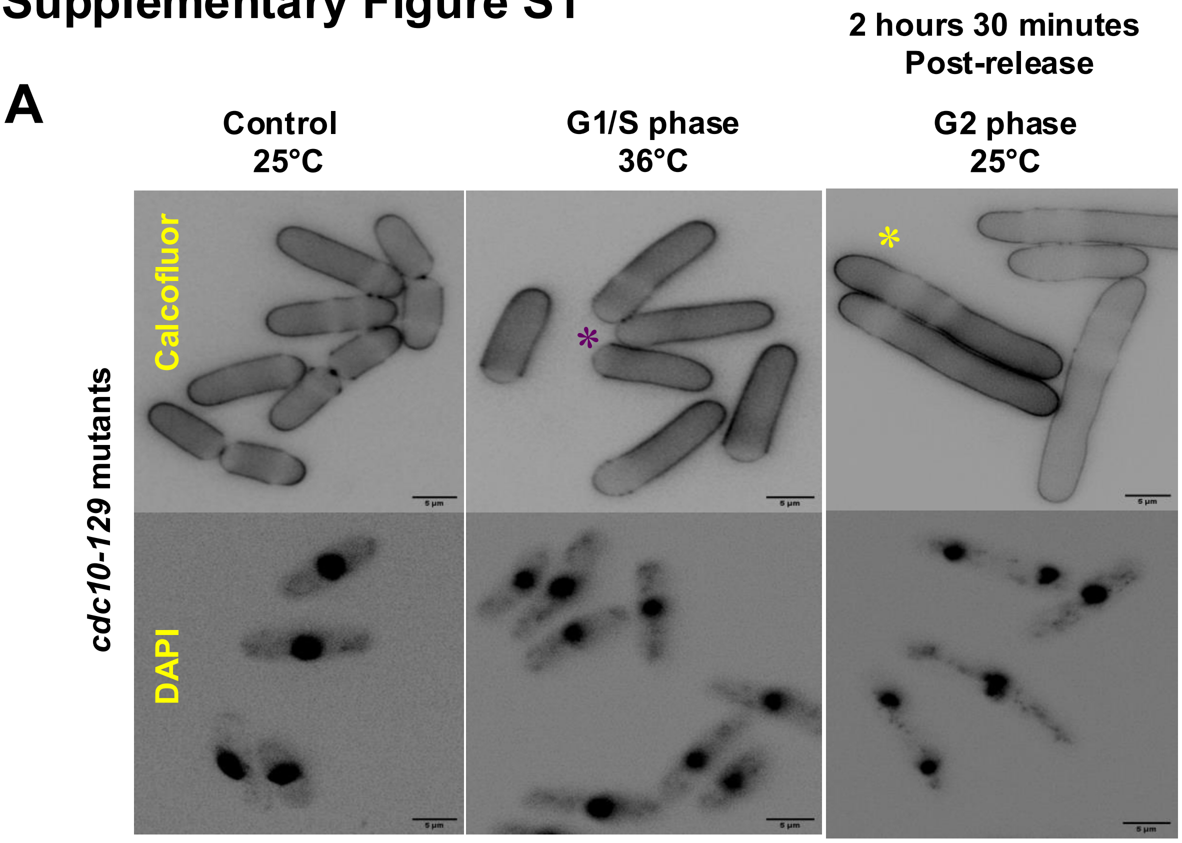


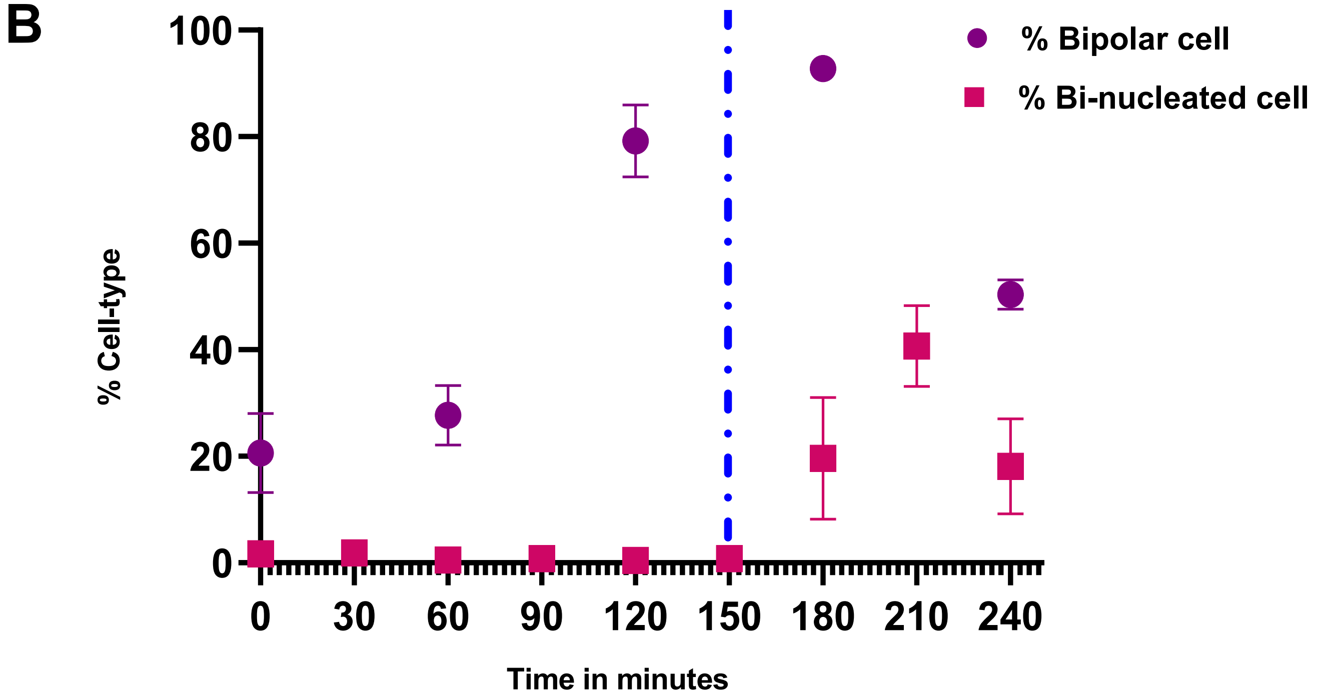


**Supplementary Figure S1: Cell-cycle progression analysis of cdc10-129 temperature sensitive mutant cells post-release at 25°C using DAPI and Calcofluor staining: A)** Calcoflour and DAPI stained cells as indicated. Purple asterisk mark monopolar cells and yellow asterisk mark bipolar cells. **B)** Quantification of Time V/S Percentage cell-type post-release at 25°C. Purple dot represents the % Bipolar cells whereas pink square represents the % Bi-nucleated cells, error bar represents the standard deviation.

#### Supplementary Figure S2

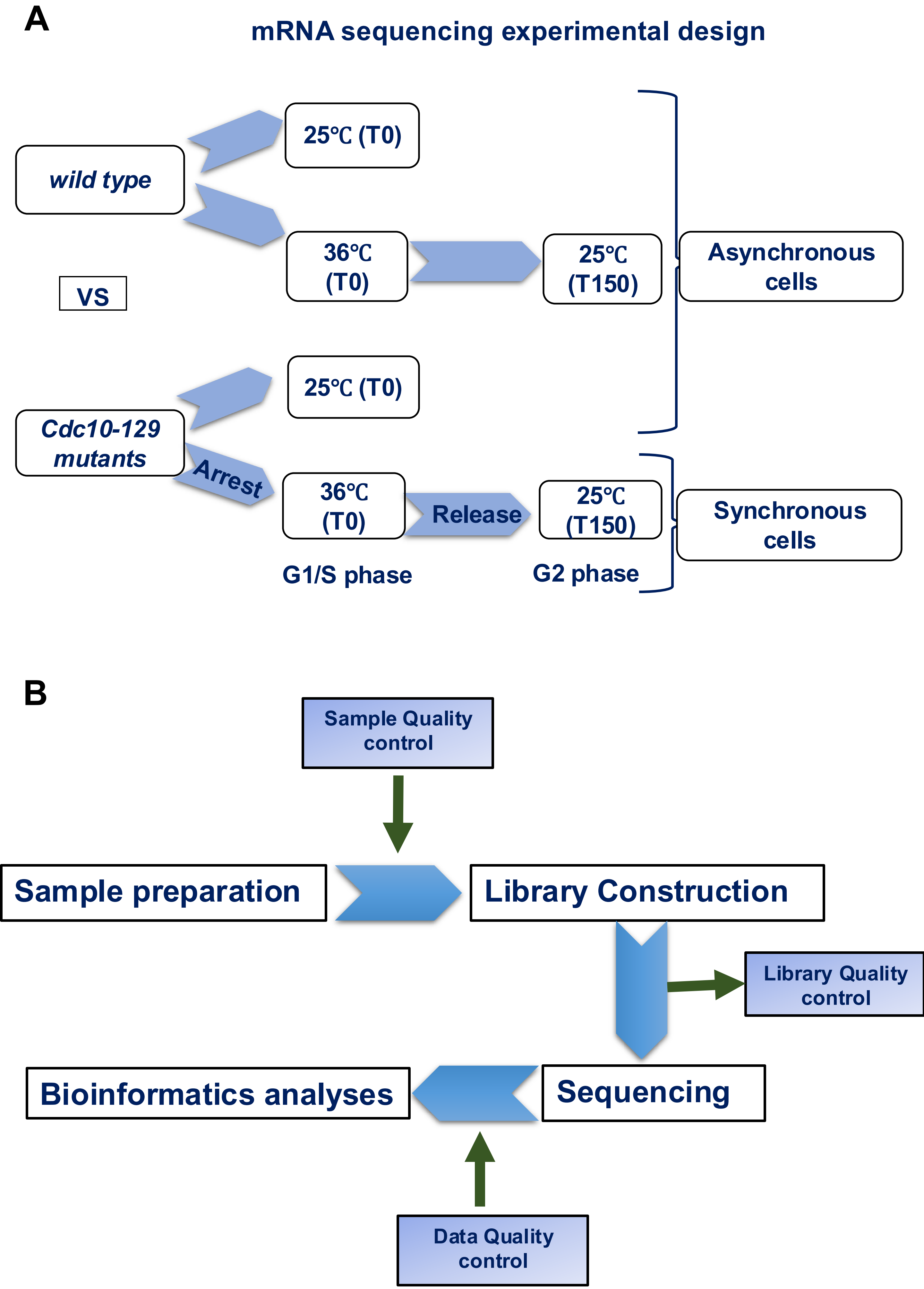

**Supplementary Figure S2: Experimental design for high-throughput mRNA sequencing and bioinformatic analysis.** A) High-throughput mRNA sequencing using *cdc10-129* synchronized cells. B) Step-wise illustration of RNA sequencing via Illumina platform based on the mechanism of sequencing by synthesis.

Supplementary Figure S3

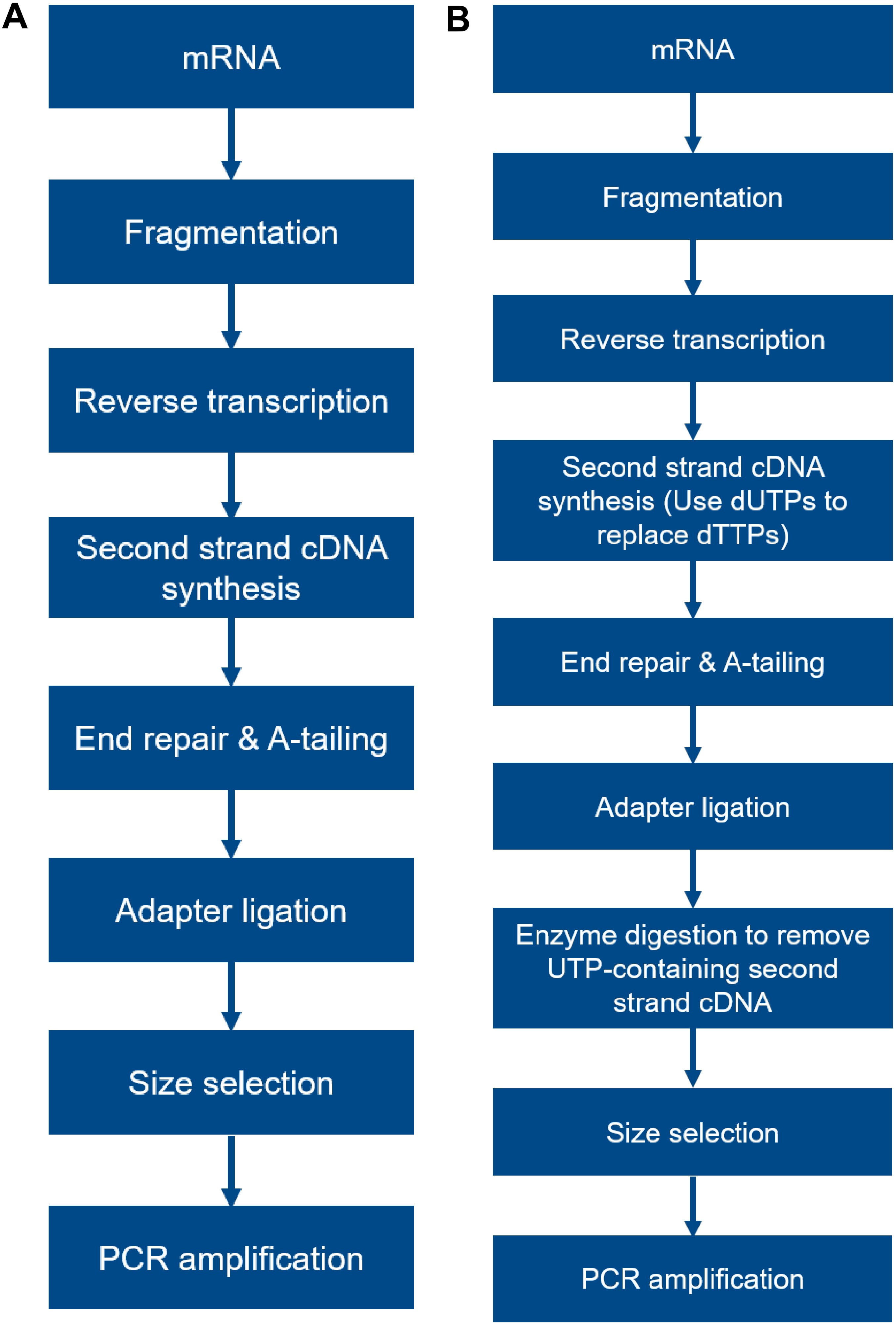

**Supplementary Figure S3: Workflow for library preparation for mRNA sequencing of *cdc10-129 cells* and wild type cells.** A) Workflow of non-directional library construction. B) Workflow for directional library construction.

### Supplementary Figure S4

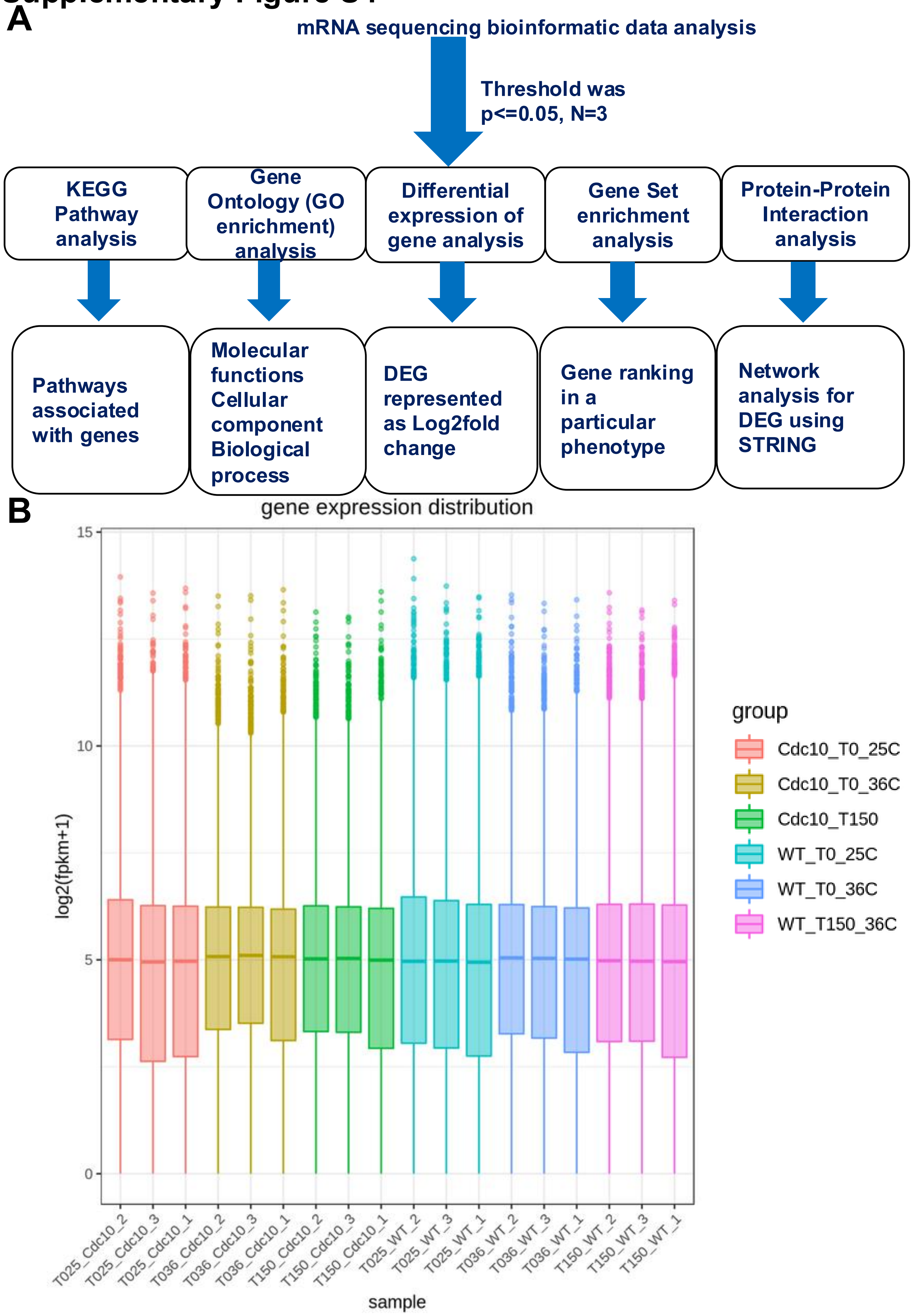

**Supplementary Figure S4: Different Analysis Method used for high-throughput mRNA sequencing data comparison between G1/S phase and G2 phase cells.** A) Data analysis for differential expression of genes, gene set from 3 independent biological replicates (N=3) with threshold of $p<0.05$; B) Sample gene expression distribution box plot. X axis represents the name of the sample, Y axis indicates the log2(FPKM+1), parameters of box plots are indicated, including maximum, upper quartile, mid-value, lower quartile and minimum.

#### Supplementary Figure S5

**A**

**B**

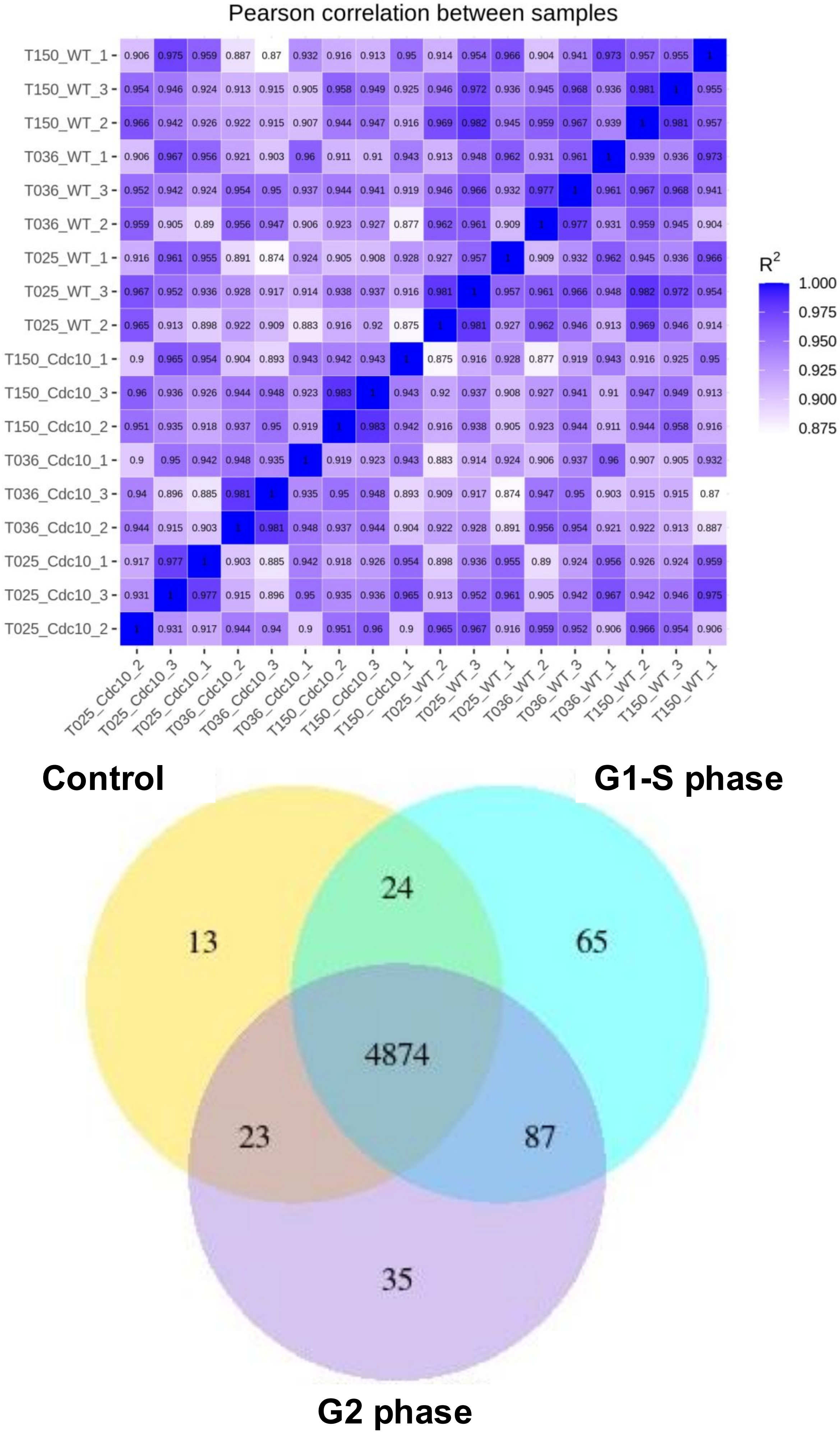

**Supplementary Figure S5: Data Quality control and analysis for mRNA sequencing.** A) Inter-Sample correlation heat Map where R2 is Square of Pearson correlation coefficient (R). B) Differential expression of gene represented as Venn diagram with control *cdc10-129* cells T0 @25°C , synchronized G1/S arrested *cdc10-129* T0 @ 36°C and synchronized G2 phase cells post-release T150 @ 25°C.

### Supplementary Figure S6

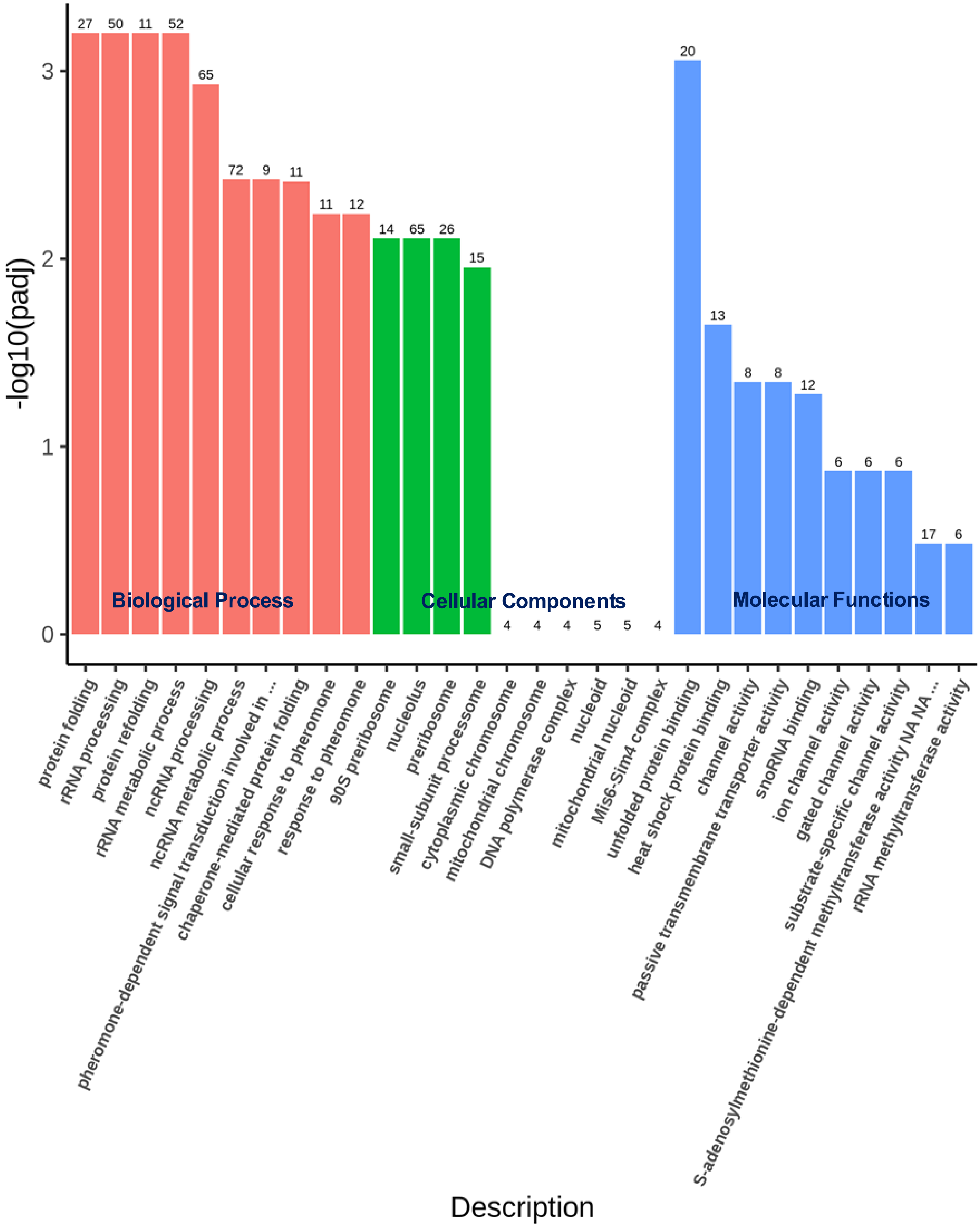


**Supplementary Figure S6: GO enrichment histograms for genes upregulated in G1/S phase cell.** The abscissa in the figure shows GO Terms, and the ordinate is the level of significance of enrichment, expressed as -log10(padj). Different colors represent different functional categories**.** Orange for Biological Process, Green for Cellular component and Blue for Molecular Functions.

### Supplementary Figure S7

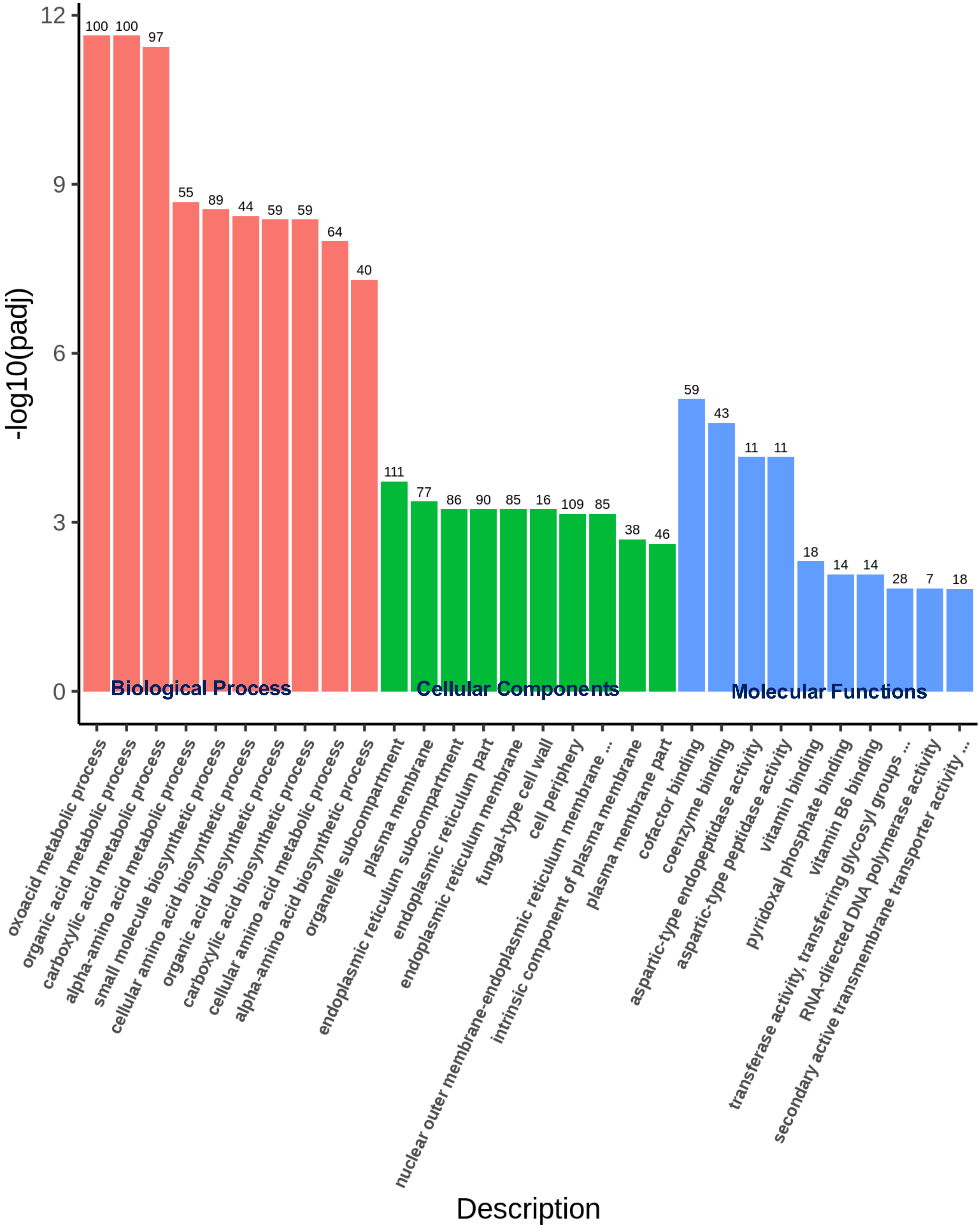


**Supplementary Figure S7: GO enrichment histograms for genes upregulated in G2 phase cell.** The abscissa in the figure shows GO Terms, and the ordinate is the level of significance of enrichment, expressed as -log10(padj). Different colors represent different functional categories**.** Orange for Biological Process, Green for Cellular component and Blue for Molecular Functions.

### Supplementary Figure S8

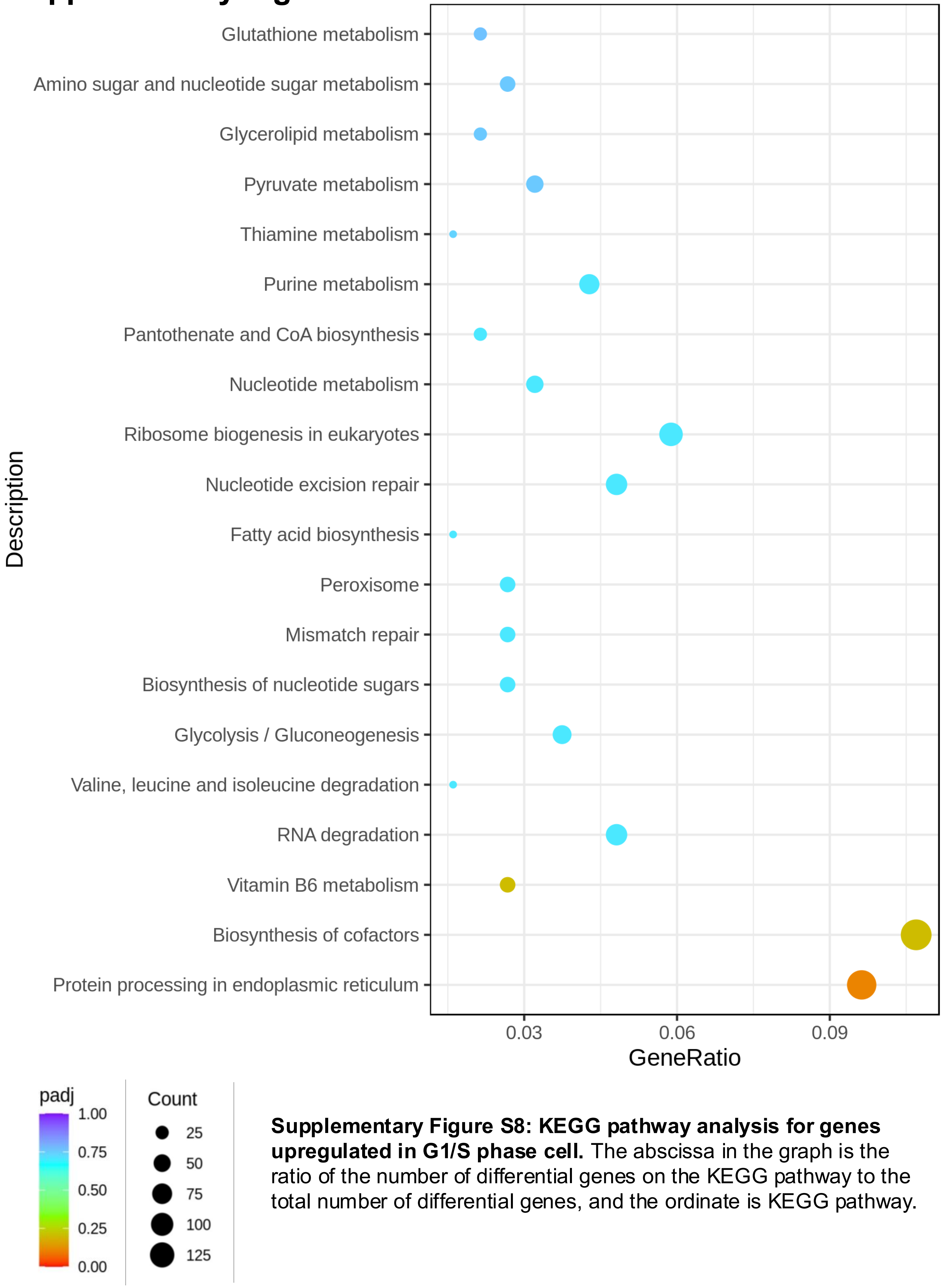


**Supplementary Figure S8: KEGG pathway analysis for genes upregulated in G1/S phase cell.** The abscissa in the graph is the ratio of the number of differential genes on the KEGG pathway to the total number of differential genes, and the ordinate is KEGG pathway.

### Supplementary Figure S9

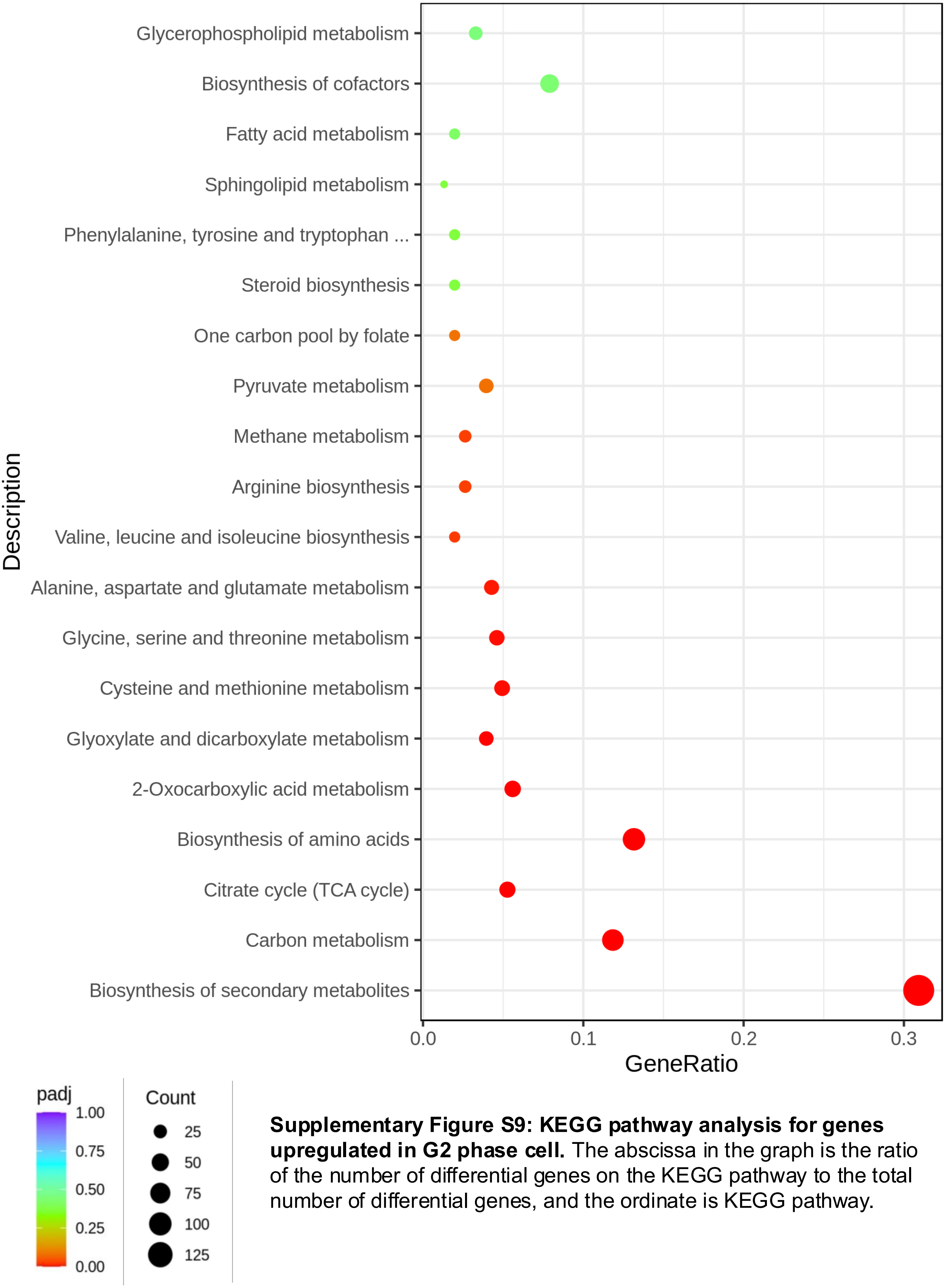


**Supplementary Figure S9: KEGG pathway analysis for genes upregulated in G2 phase cell.** The abscissa in the graph is the ratio of the number of differential genes on the KEGG pathway to the total number of differential genes, and the ordinate is KEGG pathway.

### Supplementary Figure S10

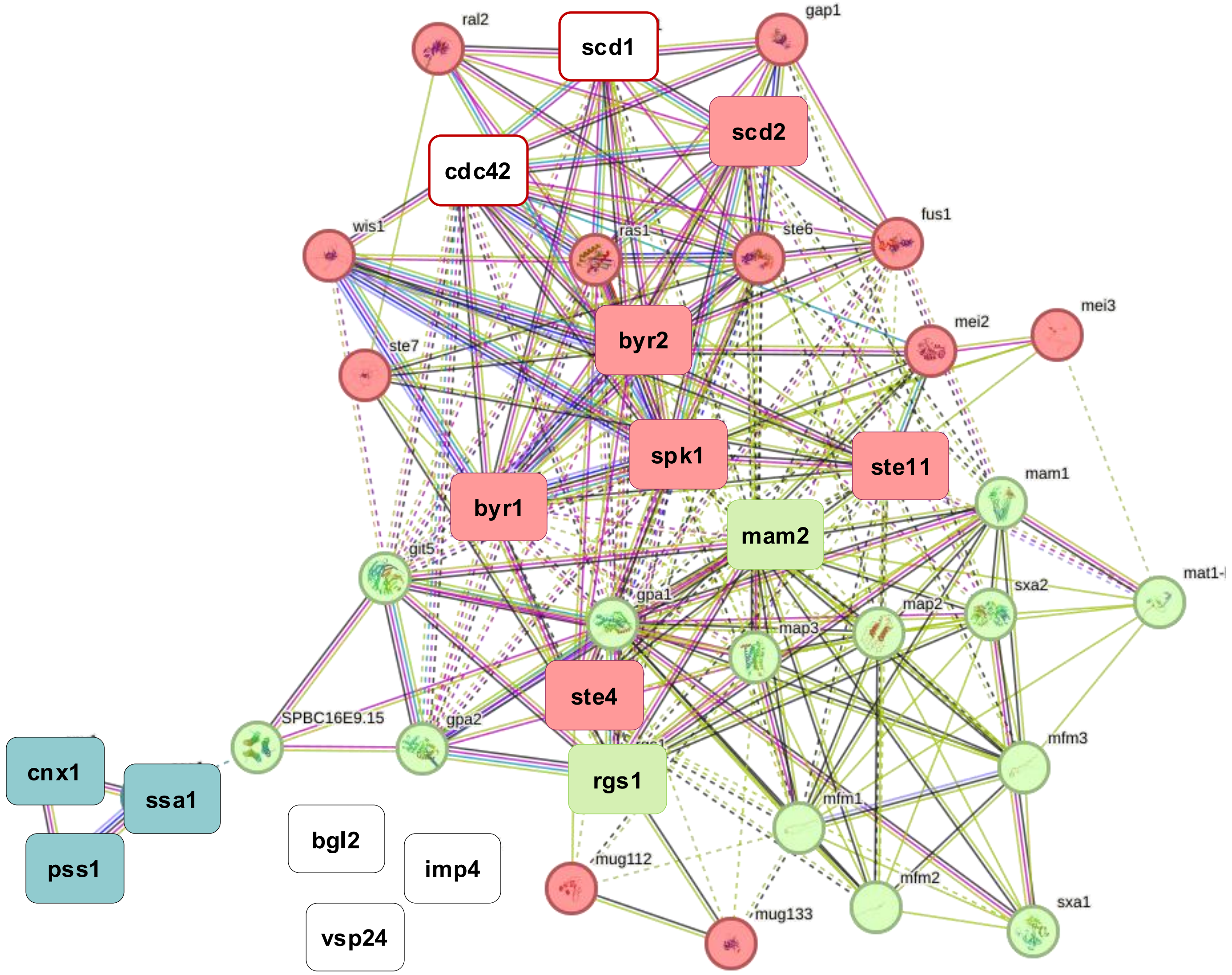


**Supplementary Figure S10: Protein-Protein interaction analysis using string for upregulated genes in the G1/S phase of *cdc10-129* cells using string.** A) Genes highlighted in magenta were pre-dominantly up-regulated in our G1/S phase monopolar cells when compared to the G2 phase bipolar cells. There are total of 40 nodes including 225 edges in the figure representing the upregulated genes in the G1/S phase cells. The average node degree is estimated to be 11.2 whereas average local clustering co-efficient is 0.628. PPI enrichment value was estimated to be < 1.0e-16. Three basic cluster appears in the PPI analysis using k-means clustering.
Red outline around the genes indicate genes involved in positive regulation of conjugation in cellular fusion. Green outline around the genes indicate genes involved in signal transduction and positive regulation of conjugation with cellular fusion and signaling receptor binding. Blue outline around the genes indicate genes involved in heat shock protein related to protein folding pathways.

### Supplementary Figure S11

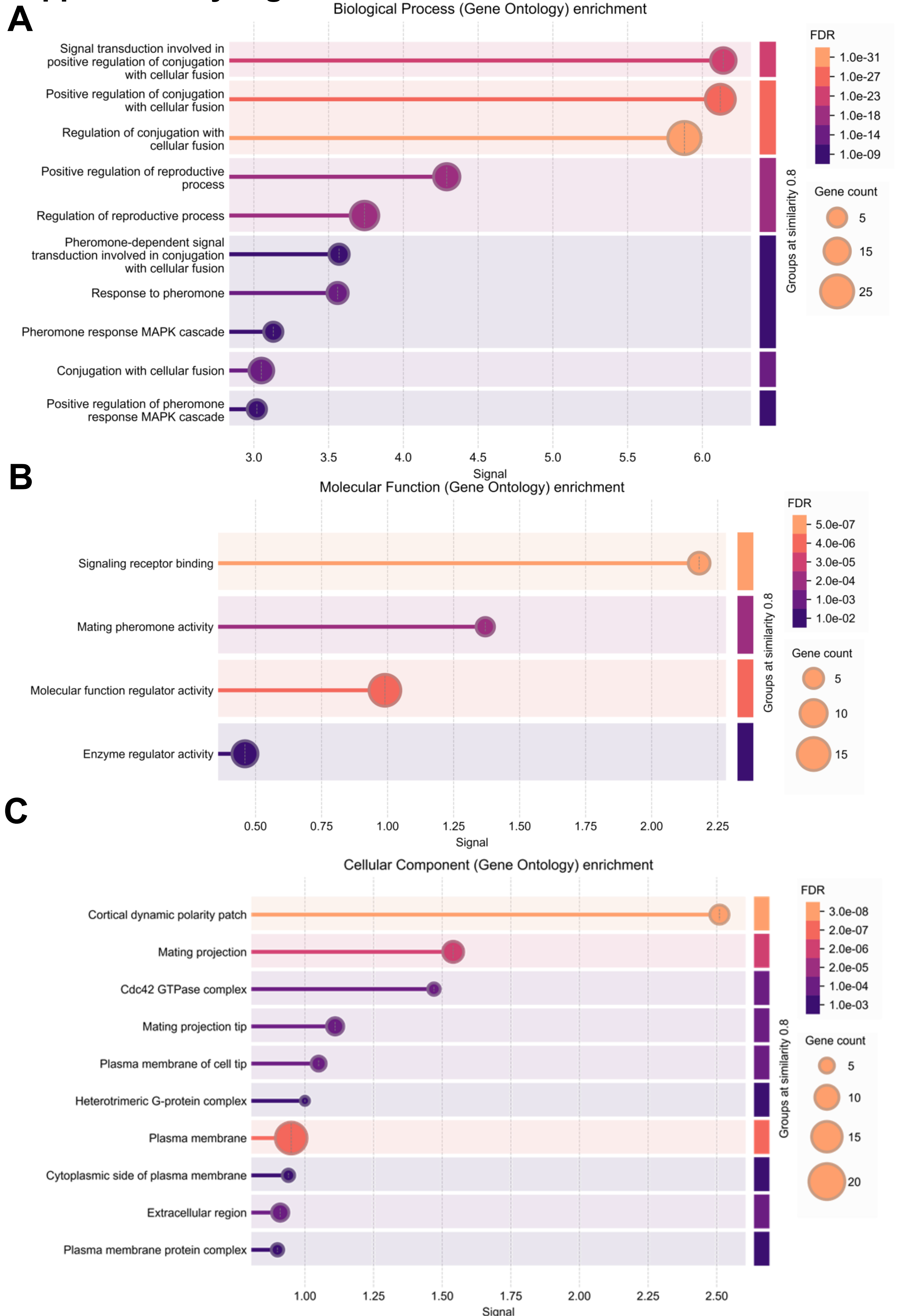

**Supplementary Figure S11: Protein-Protein interaction analysis using string for upregulated genes in the G1/S phase *cdc10-129* cells.** A) Biological Process (Gene Ontology) enrichment pathways (top 10 pathways) B) Molecular Functions (Gene Ontology) enrichment pathways (top 4 pathways) C) Cellular Component (Gene Ontology) enrichment pathways (top 10 pathways).

### Supplementary Figure S12

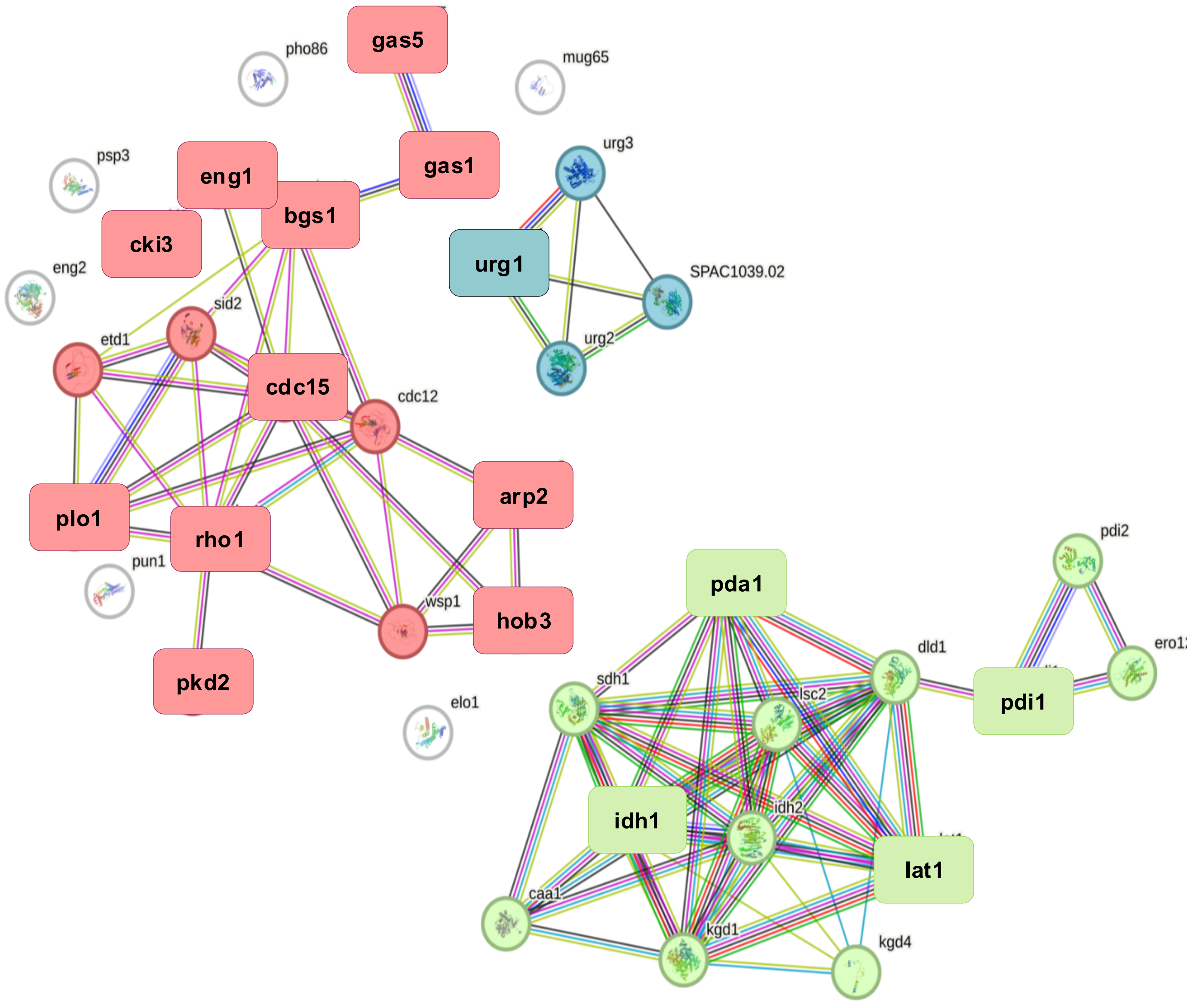


**Supplementary Figure S12: Protein-Protein interaction analysis of the downregulated gene in the G1/S phase of *cdc10-129* cells using string analysis.** A) Genes highlighted in green were pre-dominantly down-regulated in our G1/S phase cells when compared to the G2 phase cells. There are total of 38 nodes including 80 edges in the figure representing the down-regulated genes in the G1/S phase cells. The average node degree was estimated to be 4.21 whereas average local clustering co-efficient is 0.654. PPI enrichment value was estimated to be <1.0e-16. Three basic cluster appears in the PPI analysis using k-means clustering. Red outline around the genes indicate genes involved in positive regulation of cellular component biogenesis. Green outline around the genes indicate genes involved in Citric acid cycle (TCA cycle). Blue outline around the genes indicate genes involved in mixed pathways including uracil phosphoribosyltransferase, and regulation of pyrimidine-containing compound salvage.

### Supplementary Figure S13

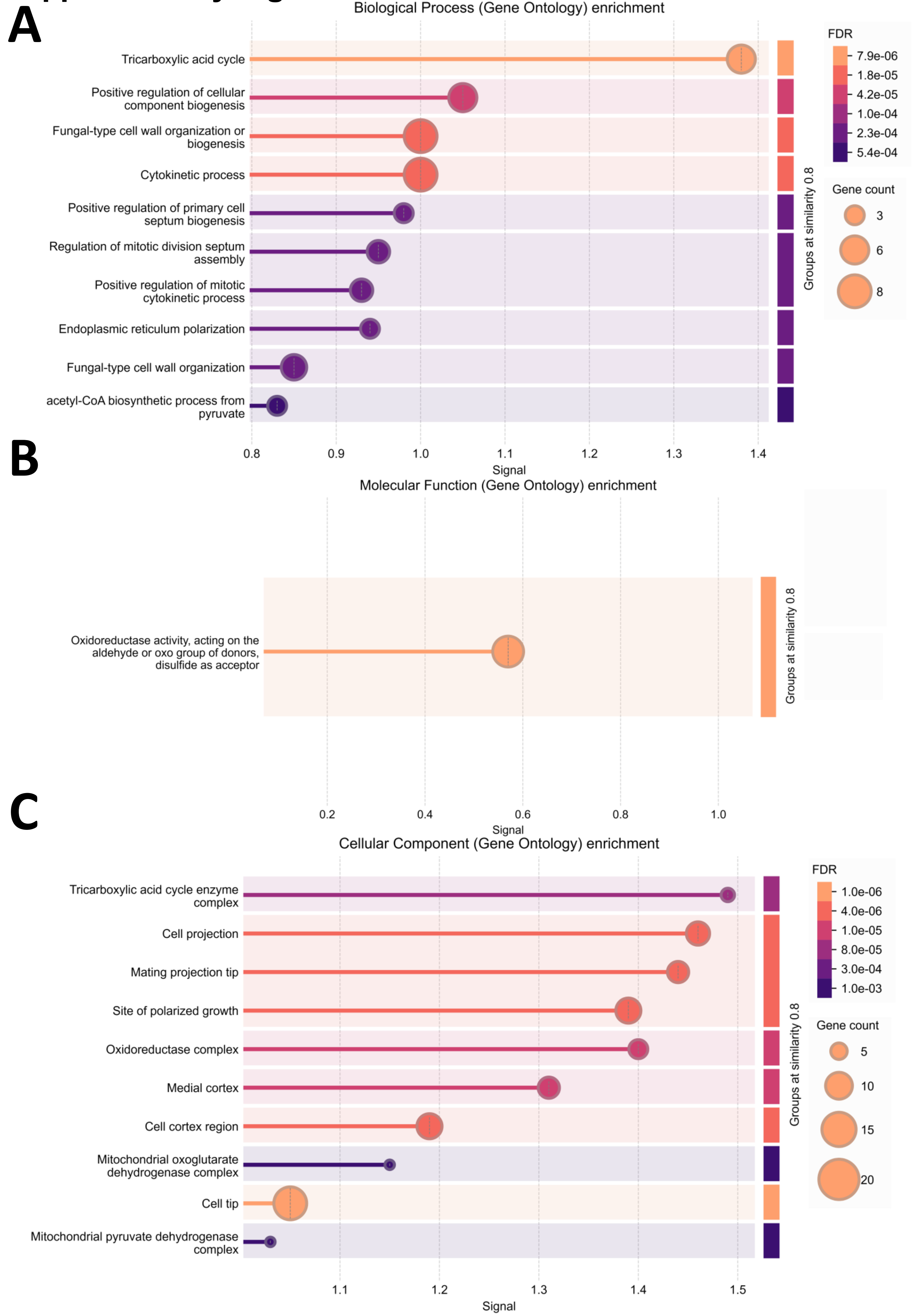

**Supplementary Figure S13: Protein-Protein interaction analysis using string for down-regulated genes in the G1/S phase *cdc10-129* cells.** A) Biological Process (Gene Ontology) enrichment pathways (top 10 pathways) B) Molecular Functions (Gene Ontology) enrichment pathways (top 1 pathways) C) Cellular Component (Gene Ontology) enrichment pathways (top 10 pathways).

### Supplementary Figure S14

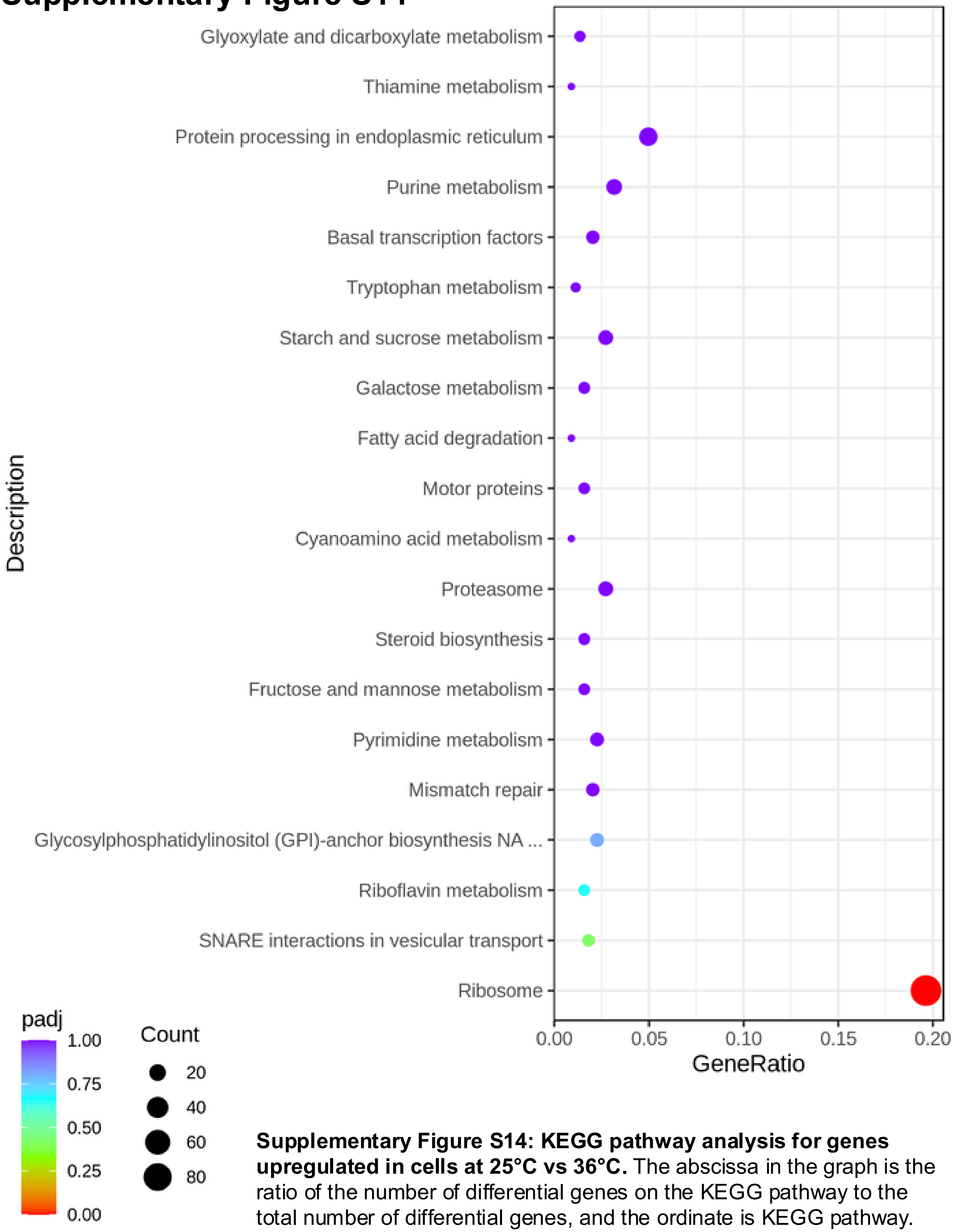


**Supplementary Figure S14: KEGG pathway analysis for genes upregulated in cells at 25°C vs 36°C.** The abscissa in the graph is the ratio of the number of differential genes on the KEGG pathway to the total number of differential genes, and the ordinate is KEGG pathway.

### Supplementary Figure S15

A

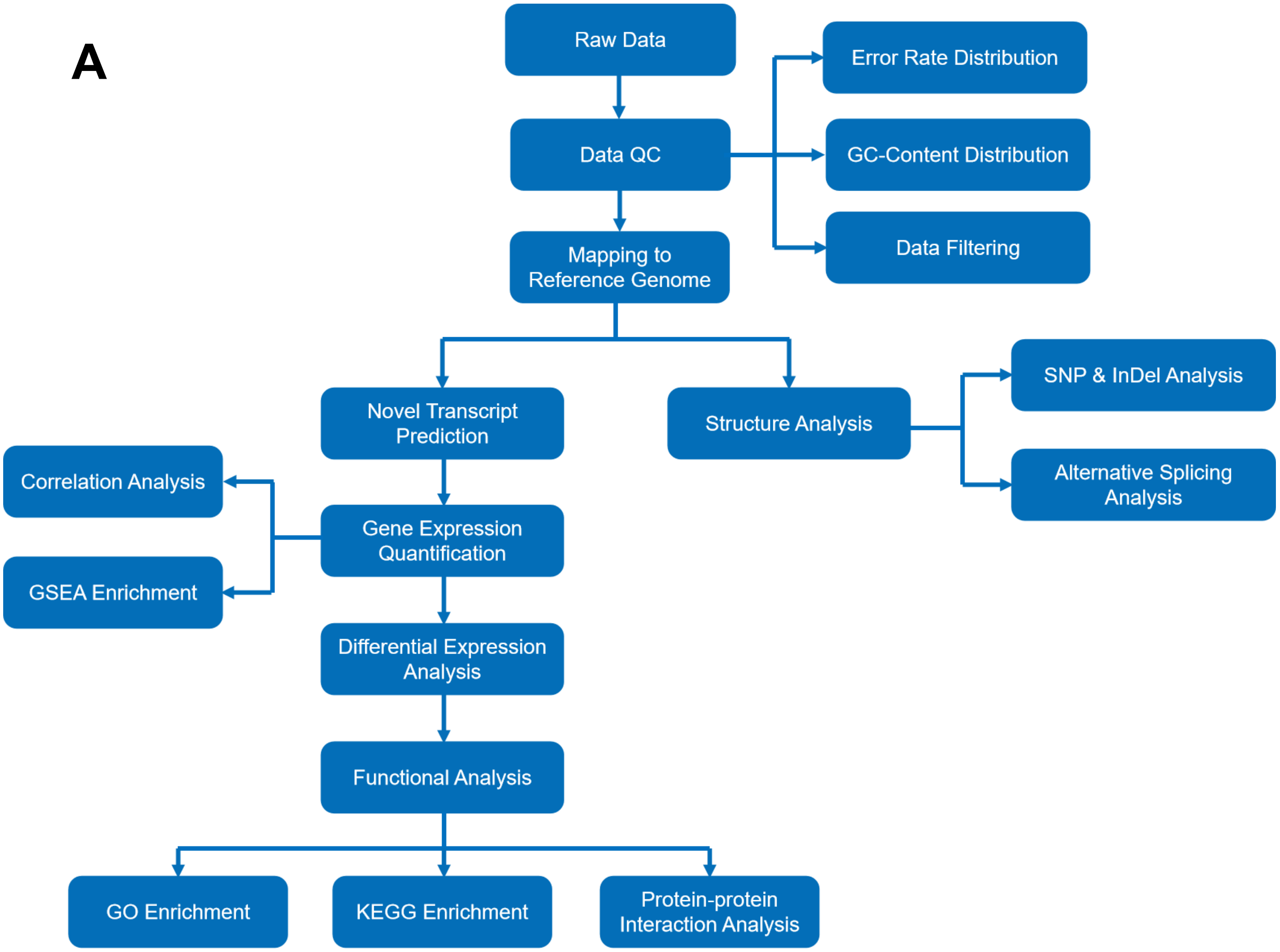


B

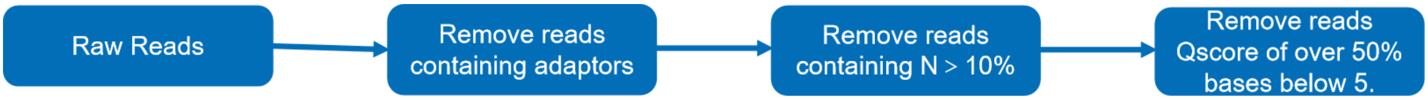


C

| Adapter | Sequence |
| --- | --- |
| **P5 Adapter** | P5-AATGATACGGCGACCACCGAGA (5’-3’) |
| | P5'-TTACTATGCCGCTGGTGGCTCT (3’-5’) |
| **P7 Adapter** | CGTATGCCGTCTTCTGCTTG-P7' (5’-3’) |
| | GCATACGGCAGAAGACGAAC-P7 (3’-5’) |

**Supplementary Figure S15: Project workflow for** A) mRNA sequencing information analysis technology flow. B) Sequencing data filtration. C) Adapter sequence for the mRNA sequencing technology used in the high-throughput raw data collection.
